# Mapping small-scale ephemeral surface water to inform transfrontier conservation planning in southern Africa

**DOI:** 10.64898/2026.04.03.715600

**Authors:** Margaret E. Swift, Anna Songhurst, Graham McCulloch, Piet Beytell, Robin Naidoo

## Abstract

Reliable freshwater access drives terrestrial wildlife movements and habitat use globally. The small, rain-fed seasonal pools critical for dryland wildlife persistence are vulnerable to rising temperatures and unstable precipitation regimes projected under climate change. In southern Africa, which is expected to warm rapidly by 2100, the drying and disappearance of surface water may cause a breakdown in seasonal migrations of large, area-sensitive, and water-dependent wildlife species. Furthermore, the disappearance of ephemeral water may concentrate wildlife around remaining surface water, increasing resource competition and human-wildlife conflict. An accurate understanding of the dynamics and drivers of seasonal surface water will therefore be crucial to wildlife and human health as climate change intensifies.

Here, we present a framework and empirical analysis of fine-scale surface water mapping in the Kavango Zambezi Transfrontier Conservation Area (KAZA), the world’s largest terrestrial conservation area (520,000km^2^). From 2019-2025, we implemented Otsu thresholding on median Automated Water Extraction Index Sentinel-2 MSI imagery (10m), creating >35 quasi-monthly KAZA-wide Ephemeral Surface Water (ESW) rasters (mean accuracy 87%, compared to 50% for a 30m Global Surface Water (GSW) product). Low wet season (November-May) rainfall drove led to low surface water fill from February-May, especially after multiple drought years (2023-2024), suggesting a critically vulnerable time for water-dependent wildlife.

Finally, we assessed the applicability of our ESW map for wildlife movement analyses. African savanna elephant (*Loxodonta africana*) are known to visit water approximately every 48 hours; a useful water map should reflect this dependence. We calculated water visitation rates using GPS data from 27 individual elephant and a threshold distance of *T =* 200m, and found that 99% of GPS tracks maintained ESW visitation rates under 48 hours, compared to only 29% of tracks for GSW visitation. These results were robust when *T* was varied, and ESW similarly outperformed random controls and several other water products, making ESW an effective proxy for actual water use in animal movement models. As climate change threatens seasonally ephemeral surface water, fine-resolution tools such as ESW will help conservation scientists and managers to understand and predict the drivers of wildlife movements at the scale of wildlife needs.

## 1 Introduction

Consistent access to surface freshwater is a vital driver of wildlife movement and distribution across the globe, from North American deserts (Morgart et al. 2005) and Chinese rainforests (Q. Zhou et al. 2011) to the Australian outback (James et al. 1999) and African savannas (Hayward and Hayward 2012; Boyers et al. 2021). Understanding the location, dynamics, and drivers of surface water availability becomes ever more critical to global wildlife conservation efforts under anthropogenic climate change, as temperatures increase and rainfall regimes destabilize (Chung and Ramanathan 2006; Core Writing Team et al. 2007; Zhao and Dai 2022; Sardans et al. 2024). This knowledge is especially crucial in southern African subtropical ecosystems, where annual-average maximum surface temperatures are expected to increase by 4-6°C (more than twice the global rate) by 2100 (Engelbrecht et al. 2015), limiting soil moisture and surface water availability through increased evapotranspiration and reduced rainfall (Zhao and Dai 2022; Yahaya et al. 2024). At the same time, these African ecosystems are also losing many long-range animal movements (Harris et al. 2009; Pringle et al. 2023), which have historically relied on seasonally fluxing vegetation and surface water to determine their timing and direction (Holdo et al. 2009; Bischof et al. 2012; Naidoo et al. 2014). Terrestrial migrations of large mammals support biogeochemical cycling (Subalusky et al. 2017), plant productivity (Holdo et al. 2007; Staver et al. 2019; Geremia et al. 2019), fire regimes (Archibald and Hempson 2016), and soil fertility and biodiversity (Geremia et al. 2025); therefore, assessing how changes in surface water availability affect wildlife movements is key not only to ongoing conservation efforts, but to broader ecosystem functioning.

Despite this urgency and impact, most current methods of mapping and tracking seasonal surface water, especially in southern Africa, lack both the spatial and temporal resolution to address questions of resource use on the scale of wildlife movements. Landsat-derived products such as the Water Observations from Space (Krause et al. 2021, Halabisky et al. 2024) and Global Surface Water (GSW) (Pekel et al. 2016) products are able to track annual recurrence of large, permanent water bodies since 1984; however, Landsat’s 30m resolution under-predicts small, ephemeral surface water (Schaffer-Smith et al. 2022) and may fail to map seasonal inundation common to southern African riparian zones. Other studies have hand-delineated water based on finer-resolution datasets such as Sentinel-2 (10m) and Google Earth imagery (1-15m) (Naidoo et al. 2020), but these efforts would be impractical for analyzing water impacts on transboundary migrations over very large scales. Some more recent, advanced methods have accurately mapped surface water systems using Simple Non-Iterative Clustering (SNIC) (Liu et al. 2024), fully automated classification trees (Huang 2018), and automatic rule-based superpixels (RBSP) (Yang et al. 2020) on Sentinel-2 and Sentinel-1 images. These accurate but computationally heavy methods rely on copious training data, high-level computational skills, and high computing power, all of which can be limited for researchers and research projects in many parts of southern Africa. To improve accessibility of research methods without sacrificing efficacy, a simple yet robust method of mapping surface water is ideal.

The lack of small-scale, seasonal surface water maps obscures accurate assessment of how water resources truly drive wildlife movement and distribution in southern Africa. Many studies rely on maps of permanent rivers or lakes (Roever et al. 2013, Abraham et al. 2019) or well-known, sometimes artificially supplied, watering holes (Chamaillé-Jammes et al. 2007a, 2007b, Smit et al. 2007) to understand water impacts on species distributions and individual movements. These approaches are generally sufficient during the dry season, when vegetation far from rivers is desiccated or inaccessible and wildlife remain close to perennial rivers and artificial waterholes (Western 1975, Cain et al. 2012), *although see* Boyers et al. 2019). In the wet season, however, heavy rains cause quick vegetation green-up and fill many small ephemeral pans, allowing animals to range far from permanent water sources (Western 1975, Fryxell et al. 1988). For example, African savanna elephants (*Loxodonta africana*) collared in 2014 in the Okavango Panhandle doubled their home range sizes from the dry season (500-650 km^2^) to the wet season (1,100-1,200 km^2^) (Makati et al. 2025), and African buffalo (*Syncerus caffer*) in northeast Namibia substantially expanded their home ranges in the wet season (Naidoo et al. 2012). Reliance on coarse maps of permanent water sources may therefore miss some crucial sources of behavioral motivation for wildlife movements, especially in the wet season. Efforts to do this in southern Africa in the past have required manual digitizing of seasonal surface water (Pozo et al. 2018) or manual selection of water index thresholds (Ji et al. 2009, Naidoo et al. 2020); these methods are infeasible for the 520,000km^2^ KAZA. If we are to gain an accurate picture of wildlife reliance on seasonally changing resources it is critical, then, to reliably map small ephemeral surface water far from permanent rivers and pools.

The mapping and analysis of small ephemeral water sources in southern Africa has recently taken major steps forward, and this paper directly builds on two key efforts in the region. A 2020 study in Bwabwata National Park, Namibia (located within the Kavango-Zambezi Transfrontier Conservation Area, KAZA TFCA) used a hand-delineated static map of 10,000 surface water pans and Normalized Difference Wetness Index (NDWI)-generated maps to analyze ephemeral water impacts on the movements of elephants and Cape buffalo (Naidoo et al. 2020). This study found that distance to surface water was an important component of both species’ movement decisions and called for a more comprehensive and dynamic map of surface water in KAZA. A 2022 follow-up study in the same region instead automatically classified surface water bimonthly from 2017-2020 with dynamic Otsu thresholding (Donchyts et al. 2016; Yang et al. 2020) on median Automated Water Extraction Index (AWEI, (Feyisa et al. 2014)) images from Sentinel-2 (Schaffer-Smith et al. 2022). AWEI has become a well-used surface water index, which has regularly outperformed other wetness indices such as NDWI and SAVI (Schaffer-Smith et al. 2022; Laonamsai et al. 2023; Sigopi et al. 2024) and on par with MNDWI (Zhai et al. 2015; Y. Zhou et al. 2017) for regions with little snow. These approaches have been crucial to mapping surface water in southern African savannas, but expanding these classification techniques to larger scales must consider variation in vegetation cover, ecosystem type, rainfall regimes, and expanding human presence.

Given these gaps and momentum, there is a need for a simple method of classifying seasonal surface water at a fine temporal and spatial resolution that is flexible enough to address landscape-scale variation in climate and vegetation. Here, we address this need by (1) mapping seasonal surface water over the largest terrestrial transfrontier conservation area in the world, KAZA, and (2) using these maps as inputs in an analysis of elephant movements in response to seasonal changes in surface water in Botswana and Namibia. We used quasi-monthly median AWEI calculated from Sentinel-2 imagery to provide seven years (2019-2025) of surface water extent and recurrence maps at 10m resolution for the entire KAZA landscape. We validated our water detection using ∼5,000 hand-selected water/nonwater points (based on very high resolution Google Earth and Planet imagery) over seven topographically and ecologically distinct areas within KAZA. We compared validation matrices for ESW to those produced for several widely used surface water mapping products: A Landsat based Global Surface Water (GSW) product (Pekel et al 2016) and three recent Sentinel-2 based products: McFeeters’ Normalized Difference Wetness Index (NDWI2), the Modified NDWI (MNDWI), and Water Index 2015 (WI2015) (Miura et al. 2025). Then, we described seasonal changes in surface water fill levels for regions connected to a river network (Seasonal Inundation Zones, SIZ) and regions not connected to a river network (hydrosheds). Given that the latter may be vulnerable to seasonal drying, we tested whether high wet season precipitation led to high hydroshed fill in the following dry season.

Finally, to highlight the value of our Ephemeral Surface Water (ESW) product for realistic wildlife movement modeling, we conducted two case studies using hourly GPS collar data from 27 African savanna elephants in northeastern Namibia and Botswana, collared from 2010-2022. For the first, we calculated average elephant drinking intervals (hours between water visits) given our 10m ESW product, other widely used products mapping surface water products (GSW, MNDWI, NDWI2, and WI2015) and two KAZA landcover maps (WWF et al. 2007; WWF Germany et al. 2021), comparing these estimated drinking rates to drinking intervals given in the literature. Elephants are well known to drink about every two days (Viljoen 1988; Leggett 2006; Naidoo et al. 2020; Pontzer et al. 2020); if a water map is to be useful for analyses of elephant movement, estimated drinking rates should match this biological reality. If our ESW product provides biologically relevant improvements in surface water mapping, we expect our case studies to show that (1) elephant movements will have a stronger correlation with surface water when using ESW than when using other surface water products to define water locations, and (2) when using ESW to calculate water abundance, elephants will be close to permanent water in the dry season, and close to ephemeral water in the wet season.

For maximum replicability and applicability to other study systems, we used only publicly available data and computational resources to create the ESW rasters. We used Sentinel-2 surface reflectance data provided by the European Space Agency; a water index derived from basic band arithmetic (AWEI); and a simple thresholding algorithm (Otsu 1979; Donchyts et al. 2016). These calculations and algorithms were run on a cloud-based programming platform (Google Earth Engine, (Gorelick et al. 2017)) free to anyone with a Google account, and all code and outputs are stored on GitHub and a public HydroShare database (Swift 2026). Elephant location data are heavily protected for conservation purposes; therefore, for our elephant water visitation case study, only the code used to run these analyses is provided. As climate change and aridification threaten dryland ecosystems around the globe, we hope that our mapping framework and case study provide an accessible guide for describing and analyzing seasonally fluxing water resources beyond our study system.

## 2 Materials and Methods

### 2.1 Study area

The Kavango-Zambezi Transfrontier Conservation Area (KAZA; Fig. 1) is the world’s largest terrestrial transfrontier conservation area, covering almost 520,000km^2^ of five southern African countries (Botswana, Angola, Zambia, Zimbabwe, and Namibia). Established by treaty in 2011, KAZA is home to a diversity of African mammals, including the largest contiguous population of African savanna elephant in the world (Thouless et al. 2016, Bussière and Potgieter 2023, KAZA Elephant Sub-Working Group 2023). The landscape of KAZA consists of mostly sparse and open bushland, woodland, grassland, cropland, and extensive wetlands, particularly of the Okavango Delta and Linyanti Swamps. Lake Kariba is a man-made lake on the Zambia-Zimbabwe border. Precipitation falls during one wet season, from November to April, ranging from 100mm mean annual rainfall (MAR) in the south to 1000mm MAR in the north (Coldrey and Turpie 2020). Because the wet season starts in November, all seasonal definitions we use include rainfall from the previous calendar year. For simplicity, we refer to wet seasons by the latter year, e.g., the season from November 2018-April 2019 is referred to as “wet season 2019”.

**Figure 1.**
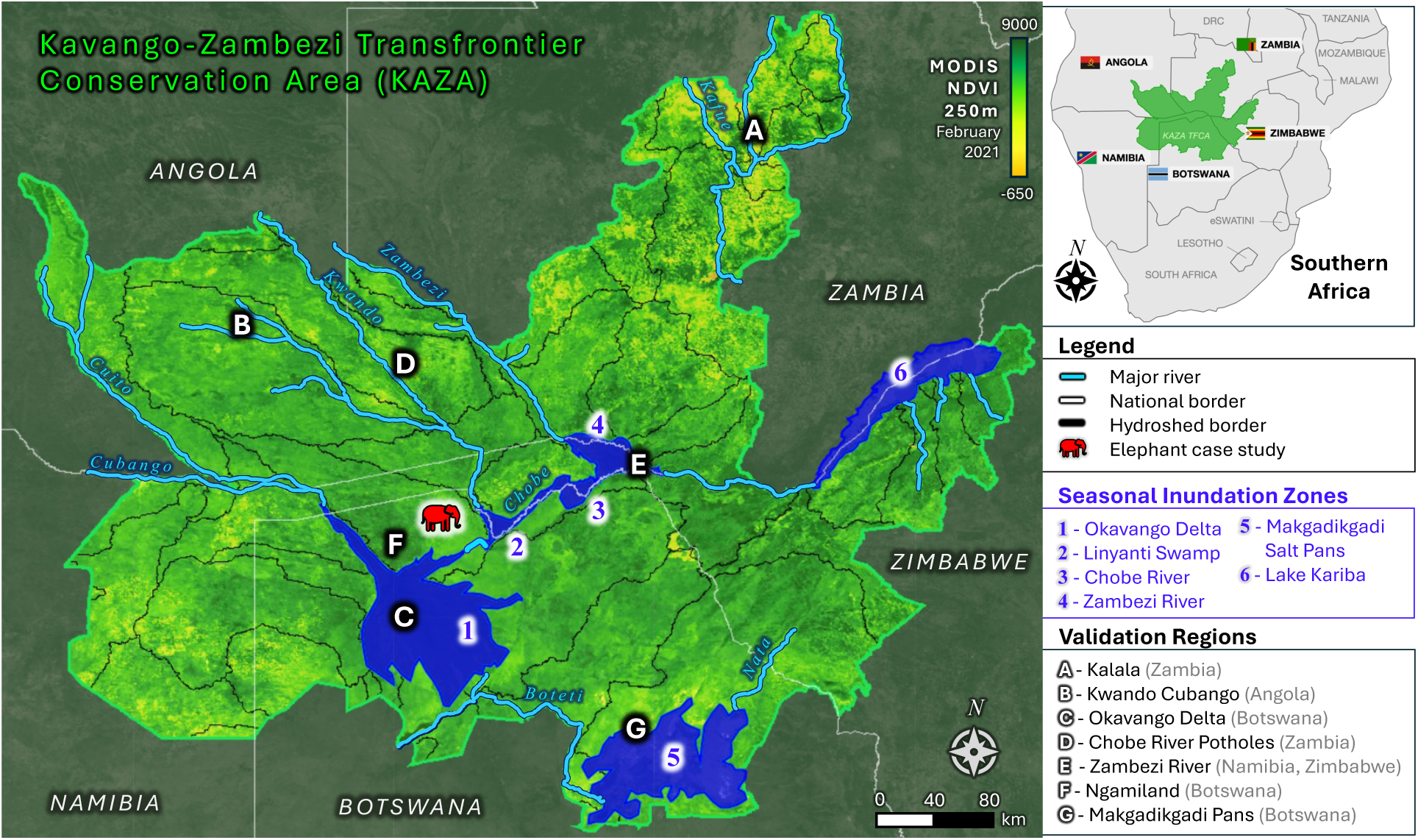
The Kavango-Zambezi Transfrontier Area (KAZA) spans 520,000 km^2^ across five countries in southern Africa (white lines). In six Seasonal Inundation Zones (SIZ; blue areas 1 - 6), water seasonally flows from rivers (major rivers in blue) and collects in deltas, swamps, and salt pans. Outside of these zones, water pools in shallow pans and is largely rainfed (divided into 54 hydrosheds, black lines). Seven validation regions (A - G) span topographical and ecological gradients. The red elephant symbol in the middle is the location of our elephant GPS case study. Background imagery for KAZA is MODIS NDVI at 250m for February 2021.

### 2.2 Geospatial data selection, coverage, and processing

While most of the KAZA hydroscape is dominated by small and shallow rainfed pools, some regions are fed by water from major perennial sources (especially the Cubango, Kwando, Kavango, and Zambezi Rivers and Lake Kariba). To address the hydrological differences between these two hydroscapes, we divided KAZA into 60 sub-regions: six hand-delineated “Seasonal Inundation Zones” (SIZ) dominated hydrologically by perennial water sources, and 54 remaining “hydrosheds” from the WWF HydroSHEDS V1.1 Level 6 dataset (Lehner et al. 2008) (Fig. 1). For this paper, a “subregion” refers to any SIZ or hydroshed; we note in the text when these subregions are treated differently. To determine the wettest years in each hydroshed, we used the Climate Hazards Group InfraRed Precipitation with Station data (CHIRPS) daily gridded (0.05°) rainfall interpolation dataset (Funk et al 2015) aggregated into mean annual rainfall (MAR) values by subregion.

SIZ were hand-selected using a combination of Sentinel-2 imagery, KAZA landcover maps (WWF et al. 2007; WWF Germany et al. 2021), and Google Maps background imagery in Google Earth Engine (GEE, (Gorelick et al. 2017)). After overlaying the “permanent water” category from both KAZA maps with Sentinel-2 imagery from March and May of 2021 (the wet season of a very wet year), we hand-delineated the maximum extent of inundation for seven SIZ in GEE. March was chosen as the period of heaviest rainfall, while May was also included to account for the timing of water drainage from the Angolan highlands into the Okavango Delta, which often leads to early dry-season peaks in inundation for the region (McCarthy et al. 2003; VanderPost et al. 2005).

Sentinel-2 MSI (10m) images were selected from December 2018 to September 2025 (seven complete precipitation years). To automate water mapping in hydrosheds, we selected Sentinel-2 imagery from five annual time periods (‘pentads’) (total *n* = 35 images). Wet season cloud coverage necessitated three-month intervals for cloud-free image selection: (1) November, December, and January; (2) February, March, and April. Dry season imagery was selected in two-month intervals: (3) May and June; (4) July and August; (5) September and October. To map the rapid changes in SIZ water levels at a fine temporal scale, we manually selected the most cloud-free monthly/ bimonthly time periods of Sentinel-2 imagery to map surface water (Appendix S1: Table S1). In this paper, a “quasi-monthly time period” refers to any of these 35 hydroshed pentads or the manually selected SIZ time periods.

All image processing was completed using GEE. To process Sentinel-2 imagery for water mapping, we masked dynamic features (clouds and burn scarring) and static features (roads, buildings, and human settlements). To mask clouds, we first excluded any Sentinel-2 image with more than 20% cloud cover. Then, we applied the “s2Cloudless” method (Zupanc 2017) with a cloud probability threshold of 20% (80% for SIZ, see Sec. 2.2) to mask individual clouds and shadows. To mask burn scars, we used the MODIS Terra/Aqua Burned Area Monthly L3 Global 500m dataset (MCD64A1), filtering on BurnDate to mask pixels burned in the two months before the Sentinel-2 image was acquired.

We masked out a number of static features that could interfere with water detection (Appendix S1: Table S2): Major road vector data were provided by OpenStreetMap (OpenStreetMap contributors 2017), then buffered by 5m on either side; settled areas were defined as a 10m binary mask of human settlements derived from Landsat 8 and Sentinel-2 (World Settlement Footprint 2015 (Deutches Zentrum für Luftund Raumfahrt (DLR) 2015)); buildings were defined as any pixel with a model confidence value > 0.5 (Google Open Buildings 2.5D Temporal Dataset, 4m resolution (Wojciech et al. 2023)), then buffered by 10m.

Data for other water mapping methods, used in validation and the elephant water visitation case study, were obtained from the following sources: GSW (Pekel et al. 2016) recurrence values; MNDWI, NDWI2, and WI2015 (Miura et al. 2025) individual rasters were downloaded and aggregated by index to gain recurrence values. In addition, KAZA05 (WWF et al. 2007) and KAZA21 (WWF Germany et al. 2021) landcover maps were downloaded and both permanent and ephemeral water classes (1, 2, and 3 for KAZA05; 80 and 81 for KAZA21) were selected to create regional water maps for the elephant water visitation case study.

### 2.3 Ephemeral Surface Water detection using median AWEI

For every pixel in each masked Sentinel-2 image in each subregion and period, we calculated an Automated Water Extraction Index (AWEI) (Eq. 2, AWEI_nsh_, (Feyisa et al. 2014)):

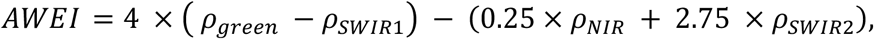

where *ρ_x_* denotes a pixel’s reflectance value for green (Band 3), Near Infrared (NIR; Band 8), Short-Wave Infrared 1610nm (SWIR1, Band 11) and 2190nm (SWIR2, Band 12) wavelengths. Pan-sharpening of SWIR bands (Wang et al. 2016) did not significantly improve the final product, and so was avoided to save time in image processing and export. We then created a median image (“median AWEI image”). We chose the median, rather than the mean, to avoid overweighting the one-time outlier values of remaining clouds, shadows, or burn scars from a single outlier image.

For each median AWEI image, we determined the optimal AWEI threshold (*AWEI\**) to distinguish water from nonwater for each subregion and time period (Fig. S2), using an Otsu thresholding algorithm in GEE (Donchyts et al. 2016). This algorithm creates a histogram of all AWEI values in a median AWEI image; this histogram is bimodal with a large peak for all nonwater and a much smaller peak for water. Otsu’s approach (Otsu 1979) finds the “valley bottom” or *AWEI\** between these two peaks by maximizing the ratio between between-class (water vs. nonwater) variance and total variance. The algorithm calculates a “sum of squares between” (SSB; a measure of between-class variance) for each of 100 histogram buckets, then returns the bucket with the largest SSB value, which is *AWEI*\*.

Otsu thresholding can fail to find a bimodal valley when there is very little water coverage, creating artificially low Otsu threshold values. We employed three methods to avoid this failure and a resultant over-classification of water: (1) We used positive masking to restrict our water mapping to only areas in which there is likely to be water (See Sec. 2.4 for a description of this method). (2) We used a minimum threshold value of −0.7 (more strict) for SIZ and −0.9 (more flexible) for hydrosheds; these values were chosen based on preliminary regional testing of average Otsu threshold values in validation regions A (Kalala), C (Zambezi), and F (Ngamiland). Finally, (3) we chose a dry-season validation period during a severe drought (July 2019), and validation regions with very little or scattered water A (Kalala), B (Kwando), and F (Ngamiland) to ensure validation coverage of low-water scenarios.

We then applied the calculated Otsu threshold to each subregional quasimonthly median AWEI image, where values below *AWEI\** were classified as “nonwater”, while those above were classified as “water”. The minimum size for ESW water bodies was 20m x 20m. This resulted in a binary raster of ESW for each subregion (SIZ and hydrosheds) and time period (hand-selected SIZ and hydroshed pentads), and exported results into a Google Cloud Storage bitbucket for further processing.

### 2.4 Hydroshed positive masking on maximum water fill

During dry times, surface water is often difficult to distinguish from background bare ground and desiccated vegetation. In addition, across a large landscape, heterogeneous rainfall can lead to massive annual and regional differences in vegetation and water conditions. We minimized false positives (i.e., avoid labeling nonwater as water) in non-SIZ hydrosheds by creating a positive “maximum water fill” (MWF) mask, restricting our final ESW maps to only areas where there was surface water in the wet season of years with the highest precipitation (Fig. 2a). This approach afforded two benefits: (1) the high contrast between vegetation and water during very wet periods allowed for easy distinction between water and nonwater pixels; and (2) because surface water in the region is generally dependent on precipitation (Schaffer-Smith et al. 2022), we captured water at its peak while minimizing Type I error.

**Figure 2.**
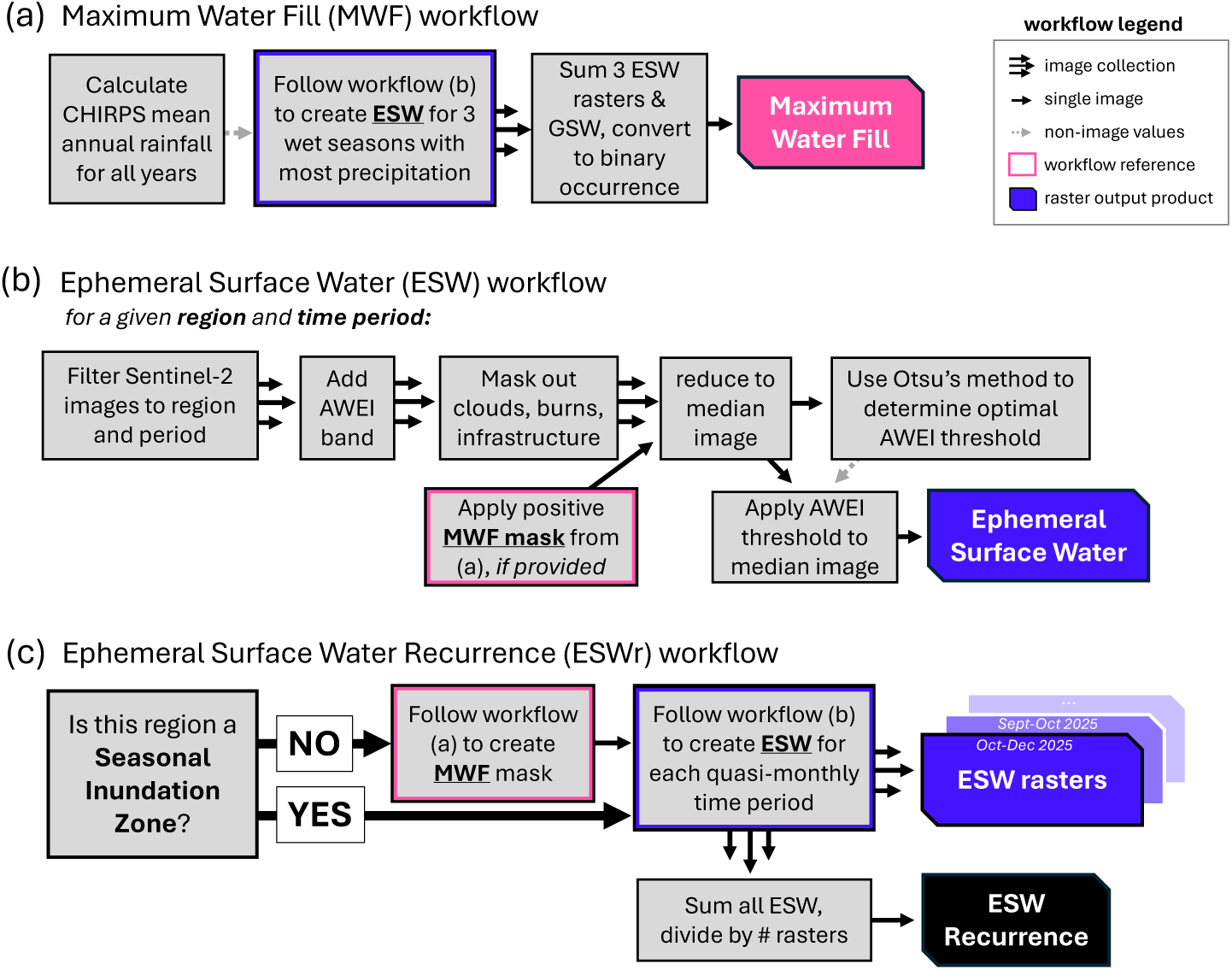
Methodological workflow for every region (hydrosheds and Seasonal Inundation Zones) in KAZA. First, if a region is not one of the six Seasonal Inundation Zones (Fig. 1, blue), a “maximum water fill” (MWF) binary mask (a) is created using the three top years by highest Mean Annual Rainfall (CHIRPS), summed with the Pekel et al 2016 Global Surface Water (GSW) product. The MWF is then used to narrow the water identification scope to only areas that had water in the wettest period, to avoid false positives. The Ephemeral Surface Water (ESW) product (b) is then created by processing Sentinel-2 images to add AWEI and mask out non-water elements (e.g., buildings, burned areas, clouds), and reduced to a single median image. An optimal AWEI threshold is calculated using Otsu’s method (Feyisa et al 2014, Schaffer-Smith et al 2022), then applied to the image to create a single ESW. For each region, all ESW are combined to create an ESW Recurrence (ESWr) raster (c), where pixel values are the proportion of images in which the pixel is classified as “water”.

To determine each hydroshed’s wettest years, we collected daily rainfall values (mm) from CHIRPS for each wet season (November 01 to May 01) using GEE, then aggregated these data to the hydroshed level to obtain the mean annual rainfall (MAR) and determine the two years with highest rainfall. We then created a MWF mask for each hydroshed by aggregating the two wettest years’ ESW Pentad 2 maps, along with the Landsat based 30m Global Surface Water (GSW) product (Pekel et al. 2016), into a single binary raster (0=never water, 1=at least once held water). Finally, we applied this positive MWF mask to all ESW rasters for each (non-SIZ) hydroshed (Fig. 2b), restricting the final maps to only areas of highest water fill.

### 2.5 Ephemeral Surface Water recurrence raster creation

Given the 35 ESW rasters for each hydroshed (with positive MWF mask applied) and the 50-60 ESW rasters for each SIZ (no MWF mask), we created two seasonal recurrence rasters: ESWr_wet_ (ESW rasters from Pentads 1 and 2, i.e., November 1 through April 30) and ESWr_dry_ (ESW rasters from Pentads 3-5, i.e., May 1 through October 31), as well as one total ESWr_all_ (ESW rasters from all time periods) (Fig. 2c). For each subregion (SIZ or hydroshed) and season (wet, dry, and total), we stacked all ESW and divided each pixel’s summed “water” classification by the number of ESW images in the raster stack to obtain a single raster where each pixel value represents a water fill frequency from 0.0 (never water) to 1.0 (always water), then mosaicked the subregion results into KAZA-wide rasters. All ESW maps were calculated in GEE, exported to a Google Cloud Storage bucket, then mosaicked using R version 4.5.1 (R Core Team 2025) before uploading to HydroShare.

### 2.6 Accuracy assessment

For assessment of method accuracy, we chose seven validation regions (Fig. 1, A-G) that represented the range of environmental and topographical variance across KAZA. Regions A (Kalala), B (Kwando Cubango), D (Chobe River potholes), and F (Ngamiland) represent the small rain-fed pools that dominate the landscape outside of the SIZ regions. Regions C (Okavango Delta), E (Zambezi River), and G (Makgadikgadi Pans) represent SIZ, with the latter also capturing seasonally dry salt flats. We completed seasonal accuracy assessments for ESW, GSW, MNDWI, NDWI2, and WI2015; KAZA landcover maps KAZA05 and KAZA21 were excluded from seasonal accuracy assessment as they are measures of general water availability.

We used a combination of Planet (∼3.4m) and Google Earth (<1m) imagery to create our validation dataset, as their high spatial and temporal resolution allowed for more accurate manual distinction between water and non-water. Point selection focused on areas most likely to confound AWEI water extraction (variable vegetation, bare ground, high topographical heterogeneity, and deep shadows) and water sources large enough to be picked up at the Sentinel-2 scale (10m). Each validation region in both the dry (July 2019) and the wet (March 2021) validation periods had at least 100 validation points for water and 100 points for nonwater, for a total of 6,400 validation points (2021 wet season: 2,792 water and 3,390 not water; 2019 dry season: 1,150 water and 3,022 not water) (Table S3). We used GEE to compare hand-verified water/nonwater classification of our validation dataset (Sec. 2.3) to those attributed from the ESW rasters for both validation periods (“dry” July 2019 and “wet” March 2021).

### 2.7 Seasonal surface water fill analysis

For our surface water analyses, a region’s fill level was the number of pixels classified as “water” in a given time period. MAR was calculated using the CHIRPS dataset (Funk et al. 2015) aggregated annually over each hydroshed. We report here regional MAR for KAZA and the total hydroshed and SIZ fill levels for KAZA over the study period. To understand how precipitation drives the amount of water available in small, ephemeral water sources, we also provide an analysis of water fill levels in non-SIZ hydrosheds. Because KAZA hydroshed area and rainfall baselines are not uniform, we used the percent of MWF or *pMWF* as a measure of hydroshed water fill for every pentad-year, and percent difference from the subregion’s average MAR or *dMAR* as a measure of precipitation anomaly for every year. Given the nonparametric distribution of *pMWF*, we compared differences in *pMWF* between pentads using a Wilcox test with Bonferroni correction (to account for multiple testing).

Because our data were zero-inflated (there were many time periods with no water fill), we fit two Generalized Linear Models (GLMs) in R for our precipitation analysis: One GLM modeled whether a hydroshed had any water at all (binomially distributed response variable), and the second GLM modeled whether precipitation significantly influenced *pMWF* (nonzero, Beta distributed response variable) (‘betareg’ package V3.2). Both models used an interaction between *pentad* and *dMAR* as the independent variables, and results are reported as marginal changes in the odds ratio (OR), as Beta distributed data require a logit link function, the log of the odds ratio *OR* = *p* / (1 − *p*).

### 2.8 Savanna elephant water use analysis

#### Elephant Data

For our case study of savanna elephant surface water usage, we used GPS data from 27 elephants collared near Botswana’s NG13 district and the Kwando River (Fig. 1, red elephant symbol; see Fig S1a for a detailed map of the region). 6 male and 6 female elephants were collared in Botswana by Ecoexist Trust (under research permit EWT 8/36/4 XVII (79)) in 2015, 2017, and 2021, and 2 male and 13 female elephants collared in Namibia in 2010, 2016, 2017, and 2020 by the Ministry of Environment, Forestry and Tourism (MEFT) (Fig. S1). Collars were programmed to record a GPS location every hour, resulting in 19,849 total GPS points in Botswana, and 124,654 total GPS points in Namibia. Transmission errors resulted in 767 data points being recorded at either four- or five-hour intervals; these observations were excluded from our case study.

#### Water Data

We compared elephant water visitation rates between ESW and GSW; MNDWI, NDWI2, and WI2015; and KAZA05 and KAZA21 (Sec. 2.2). For ESW, because our Sentinel-2 derived ESW maps only span 2019-2025, and our elephant GPS data span 2010-2021, we used the ESWr_dry_ recurrence raster as a measure of ephemeral water availability in the dry season, and ESWr_wet_ for the wet season; we then turned ESWr_dry_ and ESWr_wet_ into binary “occurrence” rasters by setting any recurrence value > 0 to 1. We also included a measure of permanent water from ESW, only including pixels with greater than 15% recurrence (ESW_15_). For all other water indices, we used the maximum extent of surface water (any pixel that was classified as “water” during any time period) for maximum comparison. Finally, we randomly generated water points across the landscape to compare all water indices to an evenly mixed control level of surface water. We created three random surface water (RSW*_n_*) datasets with *n* = 1,800 (RSW*_1800_*), *n* = 2,500 (RSW*_2500_*), and *n* = 5,000 (RSW*_5000_*) water points.

#### Water visitation comparison

A water map that adequately reflects elephant water needs should show that elephant GPS paths should approach a water hole, within some threshold distance *T*, at least once within a 48-hour period (Viljoen 1988; Leggett 2006; Naidoo et al. 2020; Pontzer et al. 2020). An elephant was considered to “visit water” if its distance from water *d_w,i_* (*w* = ESW, ESW_15_, GSW, MNDWI, NDWI2, WI2015, KAZA05, KAZA21, RSW*_1800,_* RSW*_2500,_* RSW*_5000_*) was smaller than some threshold *T*. To test elephant sensitivity to the threshold *T*, we compared *T* values from conservative to flexible: 50, 100, 200, 322, and 644m; 644m was the average hourly distance traveled by the 27 elephants. For each continuous burst of elephant movement, we initiated a “time between water visits” counter *v_w_* that increased with every hour the elephant spent away from water. To determine *v_w,i_* for each elephant GPS point *i*, we calculated the distance *d_w,i_* to the nearest pixel classified as water (using the ‘st_nn’ function from R package ‘nngeo’ (V0.4.8); with “nearest” ignoring water pixels that required crossing a veterinary fence (Fig. S1b)) then used the following to decide whether to increase or reset *v_w,i_*:

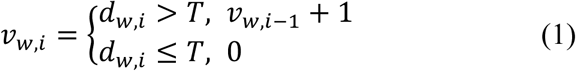

We compared threshold *T* sensitivity using a linear model for each value of *T,* with a dependent variable of distance-from-water *d_w,i_* and water type *w* as the independent variable. We used a (nonparametric) Wilcox test to determine whether the ESW visit counter *v_ESW,T_* was significantly different from *v_T_* for any other *w* for each value of *T.* For each *T,* we considered ESW to outperform other water maps if (1) the average *v_ESW,T_* was < 48 hours and was significantly different from *v_T_* for any other *w* (2) the total proportion of all data where *v_i_* < 48 was significantly greater for ESW*_T_* than for any other *w*, and (3) *d_w,i_* was higher in the wet season than the dry season for ephemeral water (ESW), and higher in the dry season than the wet season for permanent water (ESW_15_) (significant difference between means was confirmed using a *t-*test), indicating that elephant expanded their range far from permanent water in the wet season.

## 3 Results

### 3.1 Ephemeral Surface Water products

We created 1,890 ESW binary rasters for 54 hydrosheds (35 each) and 278 ESW rasters for six SIZ (Zambezi *n =* 62; Okavango *n* = 53; Makgadikgadi *n* = 71, Chobe *n* = 42, Linyanti *n* = 49, Kariba *n* = 55; see Table S2 for temporal coverage), covering the spatial extent of KAZA for seven full precipitation years (2019-2025) (Fig. 3). Optimal AWEI thresholds ranged from −0.90 to 0.02, with a mean threshold of −0.75 (± 0.11 S.D.). We also produced two seasonal rasters of ESW recurrence (ESWr_wet_ and ESWr_dry_) and one raster of overall ESW recurrence (ESWr_all_) for the study period (2019-2025), with pixel values from 0.0 (never water) to 1.0 (water in every period). Overall, the ESWr dataset provides more spatial water coverage and higher contrast between permanent and ephemeral water sources than the GSW dataset. In the Okavango Delta and Zambezi regions, for example, the difference between seasonal and permanent surface water is absent in GSW but is stark in ESWr (Fig. 3a-f); permanent river channels are clearly outlined (in pink) against regions of seasonal flooding (blue). In addition, many small water sources are completely absent in GSW but are present in ESWr (Fig 3g, h).

**Figure 3.**
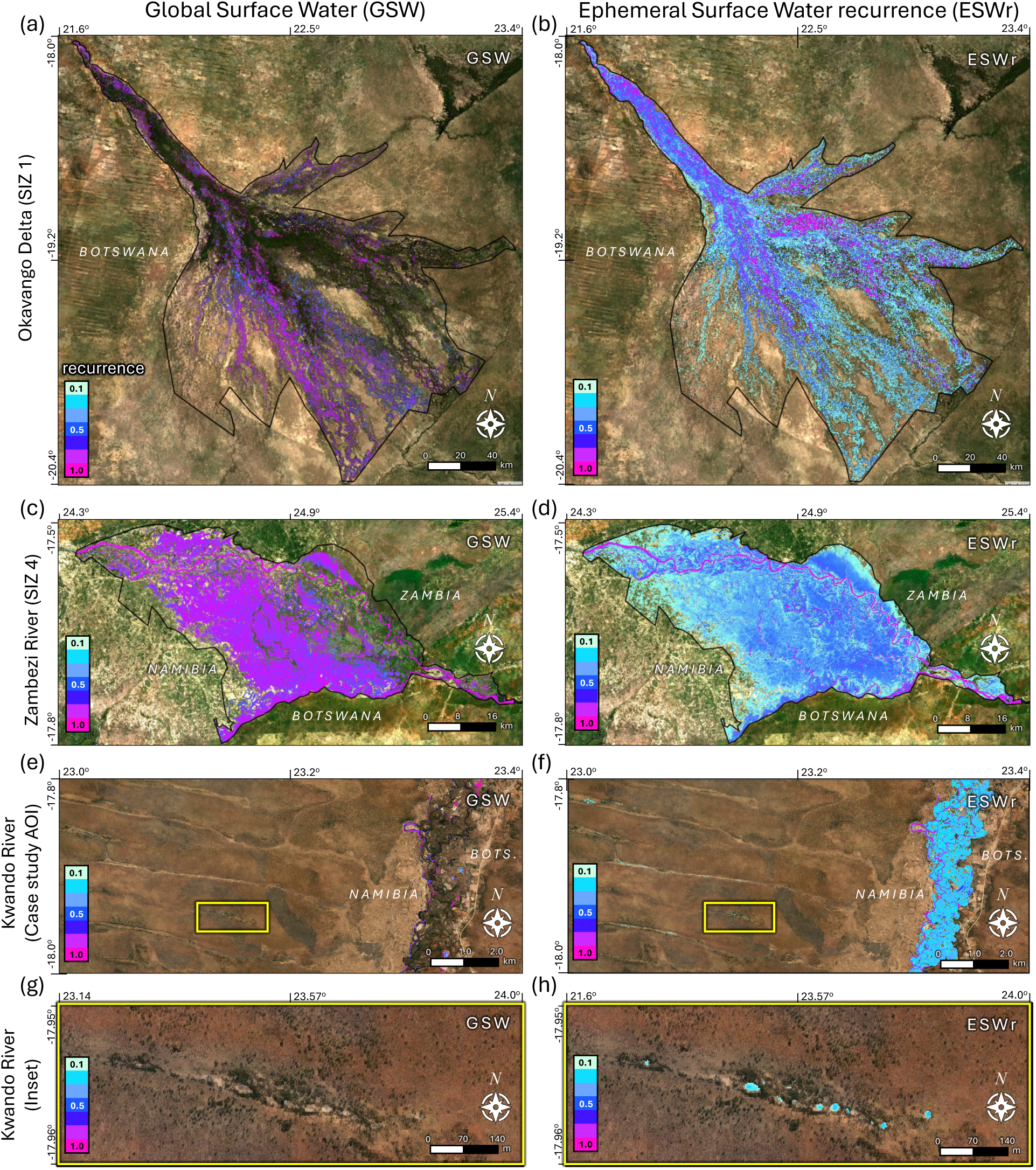
Frequency map of surface water subregions of KAZA, for both Global Surface Water (GSW) (a, c, e, g) and Ephemeral Surface Water recurrence (ESWr) (b, d, f, h). Background imagery are RGB Sentinel-2 data from June 2020. Note the increase in surface water coverage on key areas of high vegetation in the Okavango Delta (a, b); the distinct difference between the Zambezi River and its floodplain (c, d), the seasonal inundation of the Kwando River (e, f), and the omission of small ephemeral waterholes near the Kwando River (g) from GSW product (h).

### 3.2 Accuracy assessment

Our accuracy assessment found an overall vast improvement in ephemeral water mapping with ESW compared to GSW, MNDWI, NDWI2, and WI2015 (Table 1). In the wet season, we estimated a mean 92% true-water (82-99%) and 95% true-nonwater (84-100%) ESW classification compared to 51% true-water (0-88%) and 96% true-nonwater (81-100%) GSW classification. In the dry season, we estimated a mean 86% true-water (61-97%) and 100% true-nonwater (99-100%) ESW compared to 48% true-water (0-71%) and 92% true-nonwater (79-100%) GSW. Of the alternative water maps, GSW greatly outperformed MNDWI, NDWI2, and WI2015, which all performed well only in the wet season in the Makgadikgadi Pans validation region G (MNDWI: 78% true water, 79% true nonwater; NDWI2 68% true water, 84% true

**Table 1.** Accuracy assessment for the 10m ESW product presented in this paper, compared to four other surface water products (GSW, MNDWI, NDWI2, WI2015), for seven validation regions in KAZA (Fig. 1).

| Water Type | Region | ESW |  |  |  | GSW |  |  |  |
| --- | --- | --- | --- | --- | --- | --- | --- | --- | --- |
|  |  | July 2019 (dry) |  | March 2021 (wet) |  | July 2019 (dry) |  | March 2021 (wet) |  |
|  |  | *water* | *not water* | *water* | *not water* | *water* | *not water* | *water* | *not water* |
| **Scattered waterholes** | A - Kalala | **94** | **100** | **82** | **100** | **94** | **100** | **80** | **99** |
|  | B - Kwando | 61 | **99** | **87** | **98** | 0 | **100** | 4 | **100** |
| **Floodplain** | C - Okavango* | **97** | **98** | **98** | **97** | 71 | 79 | 64 | **81** |
|  | D - Chobe | **97** | **100** | **97** | **99** | 13 | **99** | 50 | **99** |
|  | E - Zambezi* | **82** | **99** | **99** | **87** | 65 | **82** | 76 | **93** |
| **Seasonally dry** | F - Ngamiland | *NA* | **100** | **87** | **100** | *NA* | **100** | 5 | **100** |
|  | G - Makgadikgadi* | *NA* | **100** | **95** | **97** | *NA* | **87** | **88** | **98** |

**Note:** Table values give the percent of “water” or “not water” validation points that were correctly classified. Cells in bold had $\geq 80\%$ classification accuracy. In July 2019, water in Ngamiland (F) and the Makgadikgadi Pans (G) completely dried up, thus their assessment was ignored (*NA*). Validation regions marked with an asterisk (\*) lie within SIZ. **Abbreviations:** Ephemeral Surface Water (ESW), Global Surface Water (GSW), Seasonal Inundation Zone (SIZ)

### 3.3 Seasonal surface water fill analysis

Surface water fill peaked in the February-May pentad for most years, tapering to a minimum in September-November (Fig. 4). For hydrosheds, the average percentage of Maximum Water Fill *pMWF* was 0.24 (± 0.30 S.D.) from November-February, 0.48 (± 0.51 S.D.) from February-May, 0.29 (± 0.32 S.D.) from May-July, 0.28 (± 0.29 S.D.) from July-September, and 0.19 (± 0.28 S.D.) from September-November. 2021 had the highest February-May surface water fill, while 2024 had the lowest (Fig. 5a), corresponding with the highest and lowest years of KAZA MAR (Fig. 4a). A pairwise Wilcox test found that *pMWF* was significantly greater in February-May (late wet season) than in all other pentads (all *p* << 0.01), and that *pMWF* was significantly greater in May-July than in July-September (*p* = 0.018) and in September-November (*p* << 0.01) (late dry season) (Table S5).

**Figure 4.**
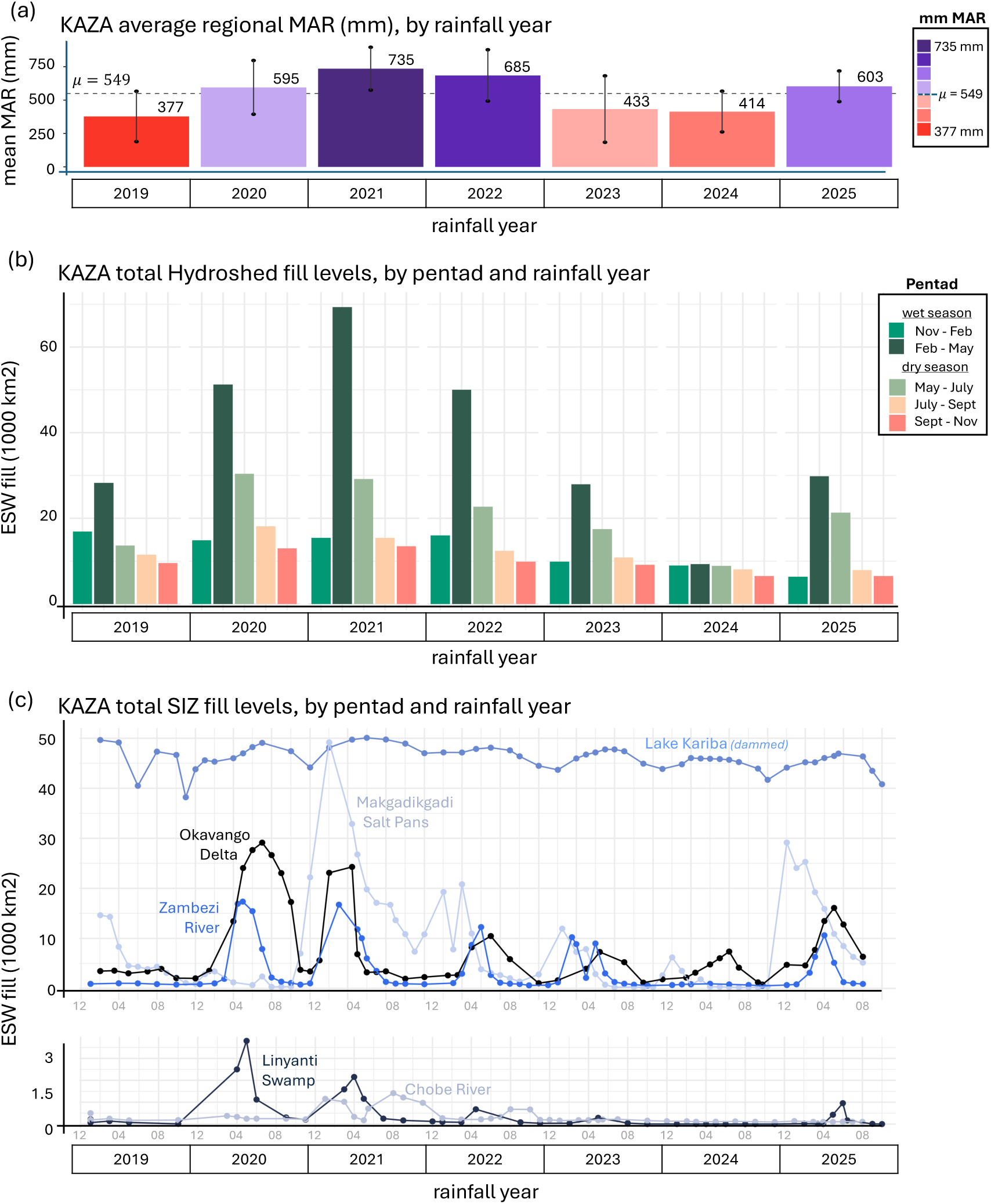
KAZA trends in mean annual rainfall (MAR) (a), hydroshed water fill levels (b), and SIZ water fill levels (c) from 2019-2025. Bars and values in (a) indicate mean MAR across all hydrosheds and SIZ, while vertical line segments indicate one S.D. above and below the mean. Bars colored purple have MAR above the KAZA mean; bars in orange are below the mean. The end of the wet season, February-May (dark green) is the period of highest fill for both hydrosheds and SIZ. Note that years of high rainfall (a) tend to correspond to years of high water fill (e.g., 2021, 2022, and 2020), especially for February-May and May-July, but that these peaks in fill are less dramatic into the dry season. Finally, 2024 was an especially low year for water fill, following two years of lower-than-average rainfall. nonwater; WI2015 75% true water, 79% true nonwater). ESW outperformed GSW by large margins in 13 cases, was almost identical in 11 cases, and slightly underperformed GSW in two cases (Table 2). We also compared accuracy tables for ESW created with and without the MWF mask, and found that ESW with MWF generally outperformed ESW without MWF, especially in the Kwando Cubango (B) and Chobe (D) regions, which are notably very dry with few waterholes (Table S4). Notably, GSW classified almost no water points correctly for the Kwando Cubango (B) or Ngamiland (F) regions (dry season 0%, wet season <10% true water).

**Figure 5.**
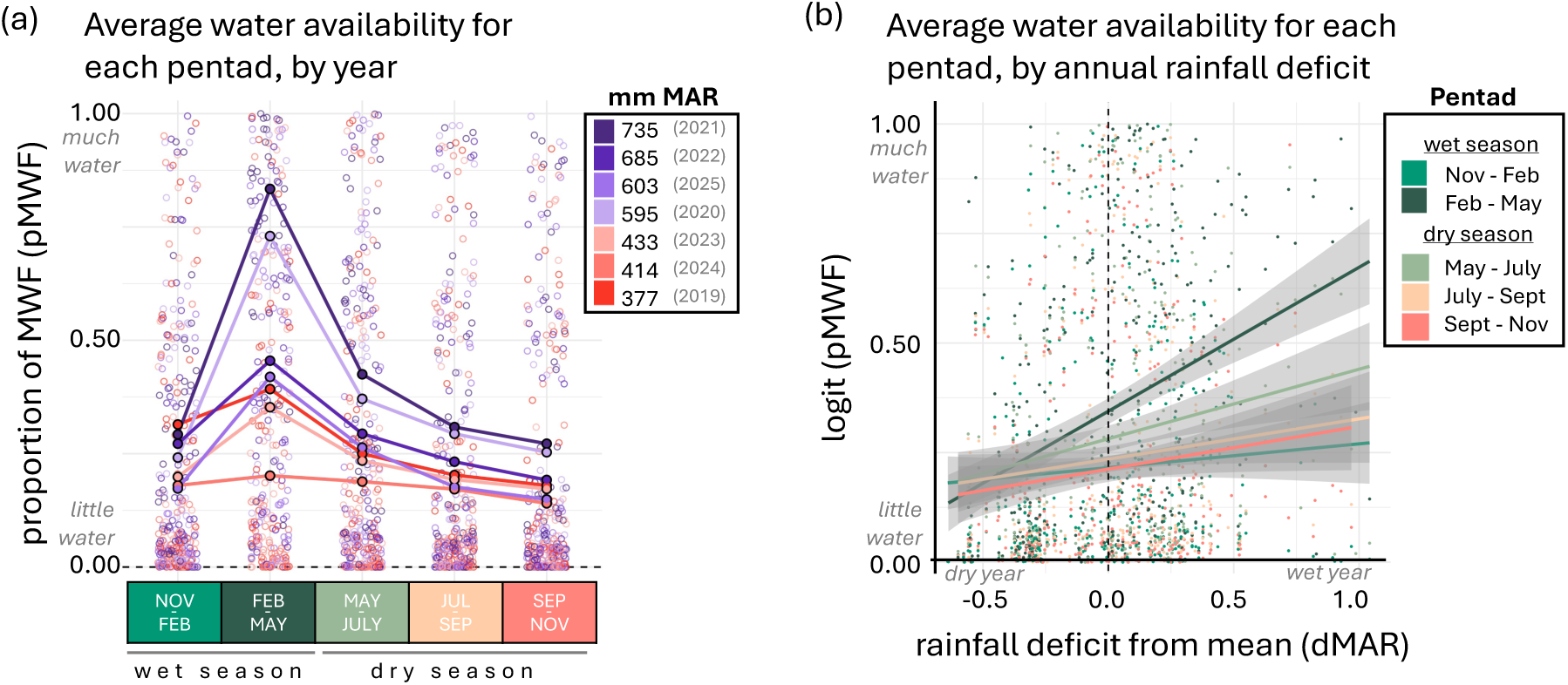
Changes in hydroshed proportion of Maximum Water Fill (*pMWF*) by pentad (a) and precipitation (b). (a) Open points represent a hydrosheds *pMWF* for a single pentad and year, while solid points and lines indicate the KAZA-wide average *pMWF* by pentad and year. At the end of the wet season, February-May, fill levels are high and the impact of high rainfall (e.g. 2021 and 2020 have high MAR and high *pMWF*; 2024 had low MAR and low *pMWF*) on the variance in fill level is evident. (b) Values given are the logit of *pMWF* for each hydroshed’s MAR anomaly (dMAR), with lines of best fit given by pentad. A Beta GLM (logit link) showed that precipitation significantly drove *pMWF* only in February-May (dark green line, *p* < 0.05), after which fill levels were unaffected by the previous year’s rainfall (all *p* >> 0.05).

**Table 2.** Surface water classification improvements of ESW over GSW, which performed best out of all alternative water maps in the accuracy assessment (Table 1).

| Water type |  | July 2019 (dry) |  | March 2021 (wet) |  |
| --- | --- | --- | --- | --- | --- |
|  |  | *water* | *not water* | *water* | *not water* |
| **Scattered waterholes** | A - Kalala | 0 | 0 | **+2** | **+1** |
|  | B - Kwando | **+61** | -1 | **+83** | -2 |
| **Floodplain** | C - Okavango* | **+26** | **+19** | **+44** | **+16** |
|  | D - Chobe | **+64** | **+1** | **+47** | 0 |
|  | E - Zambezi* | **+17** | **+17** | **+23** | -6 |
| **Seasonally dry** | F - Ngamiland | *NA* | 0 | **+82** | 0 |
|  | G - Makgadikgadi* | *NA* | **+13** | **+7** | -1 |

**Note:** Table values give the improvement in percent accuracy (ESW % – GSW %) of ESW over GSW. Cells in bold indicate an improvement of ESW over GSW (positive values). In July 2019, water in Ngamiland (F) and the Makgadikgadi Pans (G) completely dried up, thus their assessment was ignored (*NA*). Validation regions marked with an asterisk (\*) lie within SIZ. **Abbreviations:** Ephemeral Surface Water (ESW), Global Surface Water (GSW), Seasonal Inundation Zone (SIZ).

Water fill levels were significantly driven by wet-season rainfall, especially from February-May (Table 3). For our binomial GLM, we found that the OR of a hydroshed being filled (compared to the November-February baseline) was 4.85 in February-May (*p* = 0.016), and 104 with each % increase in rainfall over a hydroshed’s average MAR (*p =* 0.0044). In addition, hydrosheds had reduced OR of being filled in both July-September (OR = 0.43, *p=* 0.0041) and September-November (OR = 0.31, *p* << 0.01). For our Beta GLM, we found similar results: February-May had the highest *pMWF,* corresponding to an increased OR of 1.74 (*p* << 0.01) (compared to the November-February baseline). This effect lasted into May-July at a reduced rate (OR = 1.29, *p* = 00055). Again, *dMAR* only drove *pMWF* in February-May, with an OR of 2.8 for each % increase in rainfall over the average MAR (*p* = 0.0003) (Fig. 5b).

**Table 3.** Analysis of wet season precipitation impacts on surface water fill levels over time, as given by a binomial GLM modeling whether water was present in each hydroshed, and a beta GLM modeling the proportion of MWF filled for hydrosheds with water.

| Binomial GLM |  |  |  |  |  |  |
| --- | --- | --- | --- | --- | --- | --- |
| Variable | estimate | OR | std error | z value | p value |  |
| **(Intercept)** | 2.86 | **17.42** | 0.24 | 11.86 | << 0.01 | * |
| dMAR | 0.67 | 1.96 | 0.75 | 0.9 | 0.37 |  |
| **Feb-May** | 1.58 | **4.85** | 0.65 | 2.41 | 0.02 | * |
| May-July | -0.25 | 0.78 | 0.33 | -0.75 | 0.45 |  |
| **July-Sep** | -0.84 | **0.43** | 0.29 | -2.87 | 0 | * |
| **Sep-Nov** | -1.17 | **0.31** | 0.28 | -4.13 | << 0.01 | * |
| **dMAR x Feb-May** | 4.65 | **104.19** | 1.63 | 2.85 | 0 | * |
| dMAR x May-July | 1.44 | 4.23 | 1.02 | 1.41 | 0.16 |  |
| dMAR x July-Sep | -0.03 | 0.97 | 0.91 | -0.03 | 0.97 |  |
| dMAR x Sep-Nov | -0.18 | 0.84 | 0.88 | -0.2 | 0.84 |  |
| Beta GLM |  |  |  |  |  |  |
| Variable | estimate | OR | std error | z value | p value |  |
| **(Intercept)** | -1.06 | **0.35** | 0.07 | -15.92 | << 0.01 | * |
| dMAR | 0.18 | 1.2 | 0.2 | 0.88 | 0.38 |  |
| **Feb-May** | 0.56 | **1.74** | 0.09 | 5.95 | << 0.01 | * |
| **May-July** | 0.26 | **1.29** | 0.09 | 2.78 | 0.01 | * |
| July-Sep | 0.07 | 1.07 | 0.09 | 0.76 | 0.45 |  |
| Sep-Nov | 0.00 | 1.00 | 0.09 | -0.03 | 0.98 |  |
| **dMAR x Feb-May** | 1.03 | **2.81** | 0.29 | 3.61 | 0 | * |
| dMAR x May-July | 0.51 | 1.66 | 0.29 | 1.77 | 0.08 |  |
| dMAR x July-Sep | 0.25 | 1.29 | 0.3 | 0.85 | 0.4 |  |
| dMAR x Sep-Nov | 0.17 | 1.18 | 0.3 | 0.55 | 0.58 |  |

**Note:** Asterisks (\*) and bold values denote statistical significance. Odds Ratio (OR) = $(p / 1 - p)$ is calculated by exponentiating the fitted estimate. **Abbreviations:** Odds Ratio (OR), regional anomaly in Mean Annual Rainfall (dMAR), Maximum Water Fill (MWF).

### 3.4 Savanna elephant water use analysis

Within our case study area of interest, and within the banks of the Kwando River, our ESW raster classified 24.0km^2^ as water and GSW classified 13.34km^2^ as water. Water classification from other methods was far lower than GSW: MNDWI (65.6 km^2^), NDWI2 (6.84km^2^), WI2015 (95.9 km^2^) KAZA05 (9,887 km^2^) and KAZA21 (2,333 km^2^). Because of this lack of identified water points from other methods, our statistical analyses were only performed on ESW and GSW. For water outside of the Kwando River, our ESW method defined 1,807 individual water points ranging in size from 20m^2^ to 7,650m^2^ (mean size 354 ± 564m^2^ S.D.), approximately 6.04km^2^ of total water coverage. For this same area, the GSW product defined only 67 individual water points ranging from 20m^2^ to 2,610m^2^, (mean size 209 ± 344 m^2^ S.D.), approximately 0.13km^2^ total water coverage. On average, elephants traveled 644m (± 732m S.D.) in one hour; the results in Fig. 6 are reported using *T =* 322m, with results from other *T* values given in Table 4. Results did not significantly differ when disaggregated by sex.

**Figure 6.**
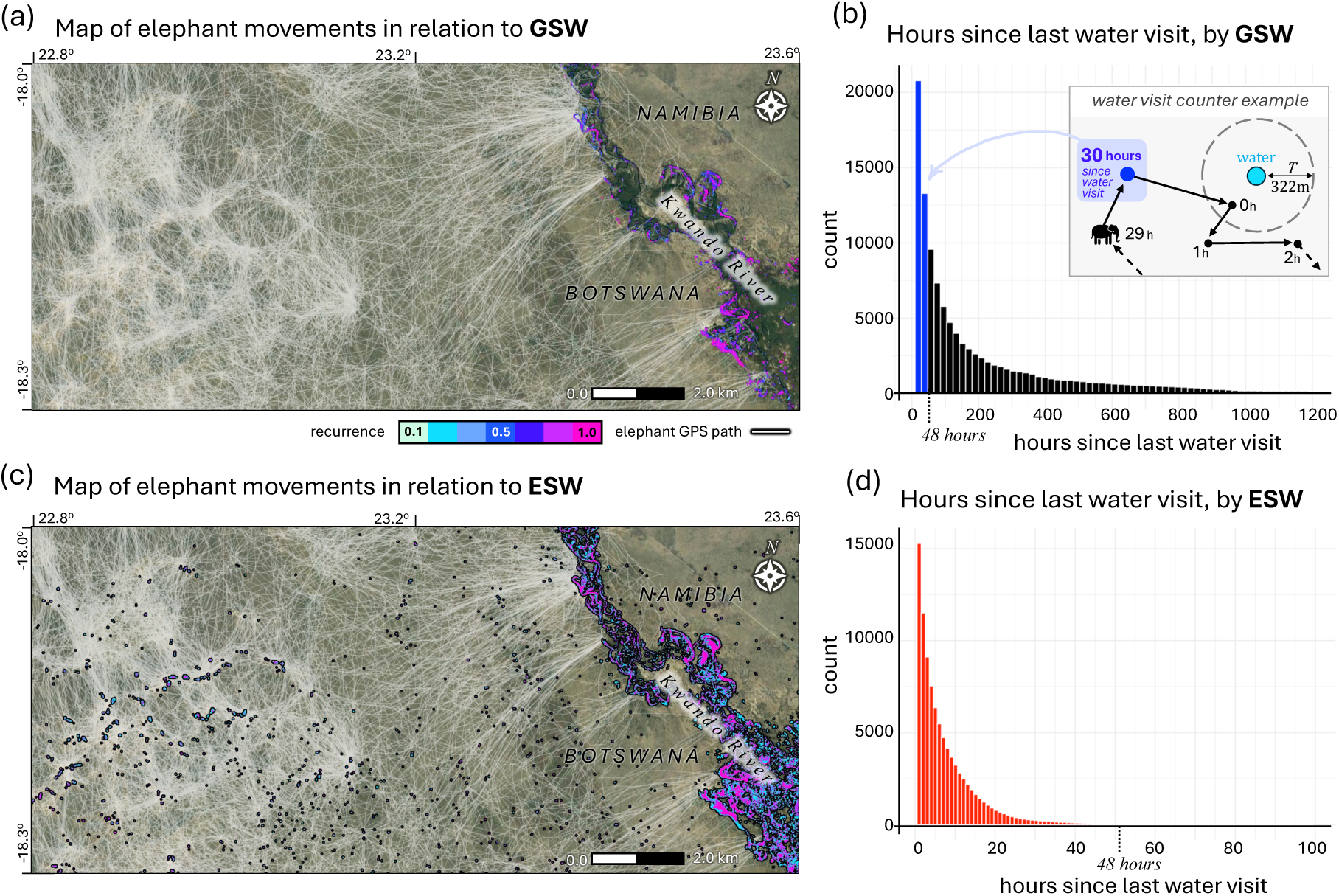
GPS collar data from 27 individual elephants from 2010-2022, mapped and summarized for GSW (a, b) and ESW (c, d). (a) When elephant GPS data (white lines) were plotted with GSW recurrence (colored dots), it was unclear why elephants clustered to the west of the Kwando River. However, a clear pattern of water dependence emerged when GPS data were plotted with ESW (c). Histograms were calculated using Eq. 1 for “hours since last water visit”, using *T =* 322m as a water visit threshold and GSW (b) and ESW (d) as water sources. See inset in (b) for a visual representation of Eq. 1. Elephants typically drink every 1-2 days; using 48 hours as a biologically relevant benchmark, we found that elephants visited GSW sources within this window only 42% of the time (b, blue bars), with a mean water visit interval of 227 (±412) hours. Elephants visited ESW sources within this window 99% of the time (d, red bars) and a mean water visit interval of 5.1 (±7.6) hours.

**Table 4.** Comparison of average time (in hours) between water visits for Ephemeral Surface Water (ESW), Global Surface Water (GSW), and Random Surface Water (*n* = 1800, 2500, and 5000 water points) products, for increasing water visitation distance thresholds (*T*).

| $T$ (m) | Average time between water visits $\bar{v}_w \pm \sigma$ hours | | | | |
| --- | --- | --- | --- | --- | --- |
|  | ESW | GSW | RSW<sub>1800</sub> | RSW<sub>2500</sub> | RSW<sub>5000</sub> |
| 50 | 27.4 ± 34.7 | 1103 ± 1127 | 309 ± 317 | 259 ± 276 | 131 ± 142 |
| 100 | **15.4 ± 18.6** | 486 ± 546 | 100 ± 122 | 73.1 ± 77.1 | 37.7 ± 42.2 |
| 200 | **8.22 ± 11.1** <sup>†</sup> | 279 ± 433 | 29.9 ± 36 | 21.9 ± 25.09 | **10.3 ± 12.5** <sup>†</sup> |
| 322 | **5.13 ± 7.63** <sup>†</sup> | 227 ± 412 | **13.55 ± 20** | **9.53 ± 12.13** | **4.15 ± 5.58** <sup>†</sup> |
| 644 | **2.01 ± 4.05** <sup>†</sup> | 181 ± 383 | **4.49 ± 12** <sup>†</sup> | **2.57 ± 4.8** <sup>†</sup> | **0.81 ± 2.1** <sup>†</sup> |

**Note:** Cells in **bold** had an average time between water visits ( $\bar{v}_w \pm \sigma$ ) less than 48 hours. Values marked (<sup>†</sup>) were not significantly different from each other, using Tukey's HSD ( $p > 0.05$ ).
**Abbreviations:** Ephemeral Surface Water (ESW), Global Surface Water (GSW), Random Surface Water of size $n$ (RSW <sub>$n$</sub> ).

The proportion of elephant GPS tracks that visited water with *T* = 322m within 48 hours was much higher when using ESW than GSW to map water. When using distance-to-GSW for water visitation classification, only *Pgsw* = 41.9% of GPS points had a time-since-last-visit value *v*_w,*i*_ ≤ 48 hours (Fig. 6a,b). When using distance-to-ESW for water visitation classification, *P_ESW_* = 99.7% of GPS points had a *v*_w,*i*_ ≤ 48 (Fig. 6c,d).

For all *T* > 50m, the average time between water visits using ESW was less than 48 hours (Table 4). Even at this strictest threshold (T = 50m), *Pesw* = 82%, significantly greater than for all RSW (*Prsw1800* = 16%, *Prsw2500* = 21%, and *Prsw5000* = 32%); ESW_15_ (21%); GSW (8%); MNDWI (9%), NDWI2 (6%), WI2015 (11%); KAZA05 (21%) or KAZA21 (22%) (Fig. 7). This stark difference holds between even ESW and ESW_15_ (ESW with recurrence over 15% of time steps), suggesting that it is the low-recurrence (ephemeral) pixels in ESW, not permanent water, that is strongly correlated with elephant movements.

**Figure 7.**
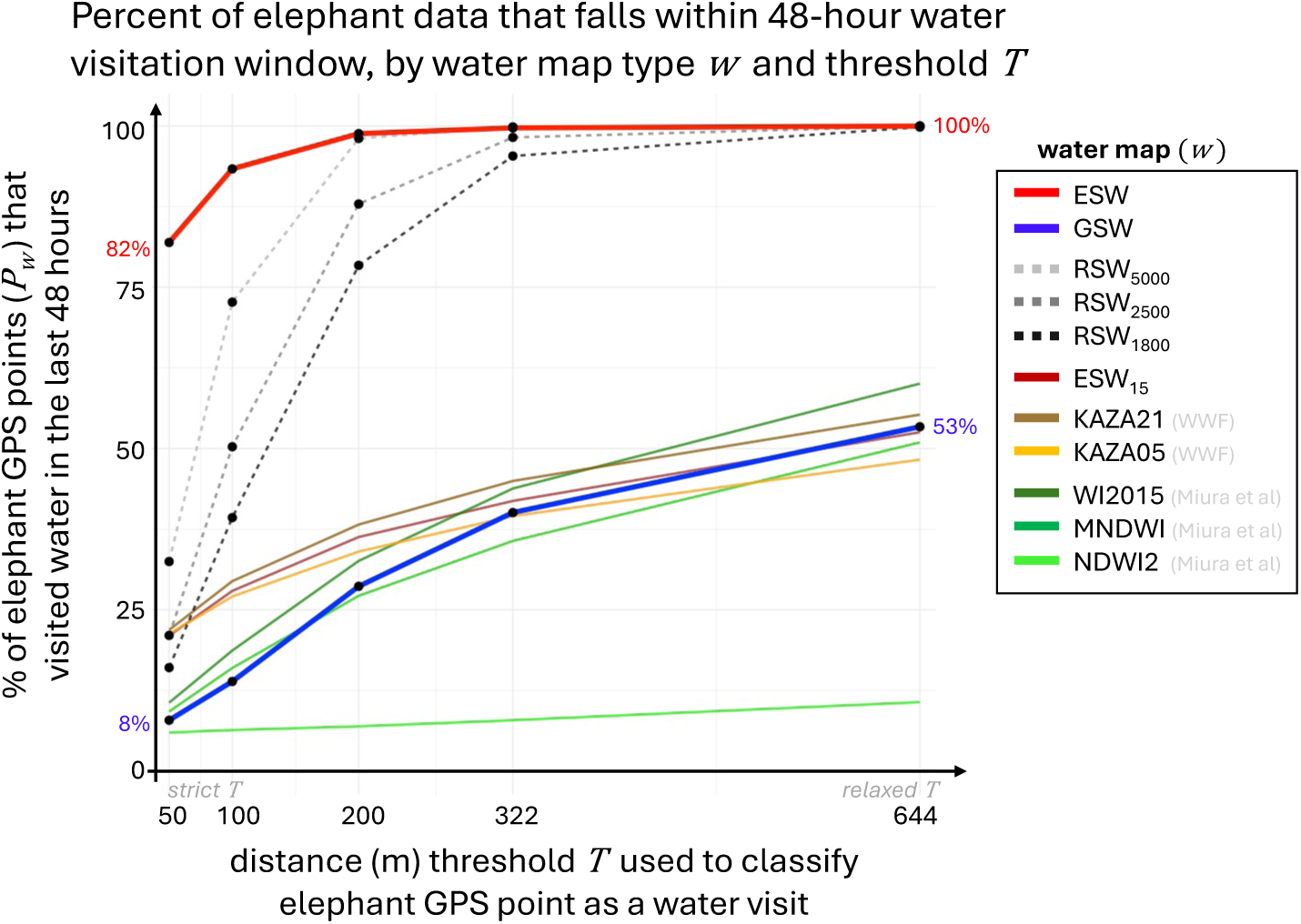
The percent of elephant GPS data with 48 hours or less between water visits (*P_w_*), given for increasing values of water visit threshold *T.* Colors indicate water maps: our Ephemeral Surface Water map (ESW, red); ESW with recurrence > 15% (ESW_15_, maroon); three sets of Random Surface Water points (RSW*_n_*, gray dashed lines), where *n* = # random points; and six other methods of mapping surface water (GSW, blue; KAZA05 and KAZA21, browns; and WI2015, MNDWI, and NDWI2, greens),. *T* = 644m is the average hourly distance traveled by elephants in this study. Even at the strictest *T* = 50m, *P_ESW_* was significantly larger than *P_w_* for all other water maps (*p* << 0.01), suggesting a strong correlation between elephant space-use and ESW. After *T =* 200m, Tukey’s HSD does not show a statistically significant difference between ESW and any RSW_n_; thereafter, ESW outperforms all real water maps but does not significantly outperform any RSW water map.

Comparing ESW, GSW, and all RSW using Tukey’s HSD, we found that all average *v^-^*_w_were significantly different from each other for *T =* 50m and 100m (*p* < 0.05); at *T* = 200m and 322m, *v^-^_ES_*_W_ was not different from *v^-^_RS_*_W5000_ (*p* < 0.05); and at *T* = 644m, *v^-^_ES_*_W_ was not different from any *v^-^_RS_*_W_*_n_*(*p* < 0.05), but GSW still lagged behind. Even using the least strict threshold (*T* = 644m), *v^-^_GS_*_W_ = 181 ± 383 hours, with only 53% of *v_GSW,i_* < 48 hours (Fig. 7).

Finally, we found that elephants’ distance from ephemeral water (*d_ESW_*) was lower in the wet season (621m +/- 446m S.D.) than the dry season (597 +/- 429 S.D.) (*p* << 0.05), and that elephant distance from permanent water (*d_ESW15_*) was lower in the dry season (4538m +/- 2860 S.D.) than the wet season (5277m +/- 3203m S.D.) (*p* << 0.05), indicating a shift in elephant space-use from close to permanent water ESW_15_ in the dry season, and close to ephemeral water ESW in the wet season.

## 4 Discussion

This paper presents comprehensive 10m-resolution surface water maps for KAZA, a significant improvement in both the availability of water mapping in a rapidly drying region and in the methods available to researchers with limited computational skills or training data. While global Landsat based products such as the Global Surface Water product (Pekel et al. 2016) and global Sentinel-2 based products like MNDWI, NDWI2, and WI2015 (Miura et al. 2025) are useful for mapping persistent large water bodies over a long time and at a global scale, and regional landcover products (WWF et al. 2007; WWF Germany et al. 2021) at a regional scale, our Ephemeral Surface Water products map water at fine spatial and temporal scales. These maps focus on small, ephemeral water features that are often ignored in both global and regional products. In addition, while object-based approaches (Liu et al. 2024) and machine learning techniques (Huang 2018) are more accurate on a small scale, these methods require huge amounts of training data and high-level computational skills which may not be available for all researchers and research projects. Our relatively simple Otsu thresholding method, using open source data (Sentinel-2 MSI) and tools (Google Earth Engine, R), is accurate and approachable at many scales, filling a critical gap in the research tools available for water monitoring, wildlife conservation, and climate change research.

The ephemeral surface water recurrence maps (ESWr) we present here provide a nuanced seasonal understanding of surface water available on the landscape. For example, regions like the Okavango Delta (SIZ 1, Fig. 3a,b) and Zambezi River floodplain (SIZ 4, Fig. 3c,d) are poorly represented in global surface water datasets, and where there is water, recurrence is homogeneous across the region (Fig. 3a,c). Our ESWr product reveals much more variation in surface water recurrence than has been previously mapped, and especially highlights persistent river channels (dark pink) that are difficult to extract from single images or with the naked eye (Fig. 3b,d). Further, our seasonal analysis shows that surface water fill level significantly peaks at the end of the wet season (February-May), and that these peaks are driven by each wet season’s rainfall (Fig. 4, Fig. 5). The influence of rainfall on water fill levels does not continue after the end of the wet season, however; high wet-season precipitation does not translate to overall higher water availability for the rest of the year. In addition, although MAR was not much lower for 2024 than 2023, both hydrosheds and SIZ saw a large dip in surface water fill during 2024 (Fig. 4), which could indicate a compounding effect of two very low rainfall years in a row. As southern Africa warms and dries over the coming decades (Engelbrecht et al. 2015), shifting rainfall regimes are most likely to impact surface water, and therefore wildlife behavior, during the February-May window in which waterholes are most sensitive to rainfall.

Small ephemeral waterholes are clearly important for wildlife use and movement (Fig. 6, Fig. 7), yet are entirely omitted by global surface water products (Fig. 3g,h). Our ESW maps provide significant improvement on existing global surface water maps for all validation regions (Table 1, Table 2), and our sensitivity analysis shows a remarkable fidelity of elephants to ESW, even at the strictest threshold of water visitation classification (*T =* 50m). In addition, even within our ESW maps, a distinction lies between ephemeral (ESW) and permanent (ESW15) surface water; elephant clearly rely on permanent water in the dry season and move towards ephemeral water in the wet season (Sec. 3.4), and elephant water visitation rates at the strictest threshold *T =* 50m for ESW15 were much closer to the low visitation rates for GSW and other indices of permanent water (Fig. 7). It is clear that these small ephemeral water sources are critical for wildlife movements, and that these behaviors cannot be properly assessed with coarser data products. This case study also support use of these water recurrence maps beyond the 2019-2025 window in which they were created, and could be extended even further into the past by leveraging positive masking on Landsat data where water sources are larger than 30m.

The process of creating these water maps has highlighted some important gaps in current applications of Sentinel-2 imagery for water mapping. First, burn scarring on the landscape created significant bottlenecks in the time periods we could map water. Burned areas often appear to sensors to be similar to water, but we found that current Sentinel-2 based methods of burn masking over-corrected and removed small pools of surface water. We instead used a MODIS-based burned area map, as MODIS provides thermal bands for focusing burn scar mapping, but the 250m resolution of MODIS meant burn scar masking was coarse and often inaccurate on the margins. In addition to burn scarring, the cloud masking available directly with Sentinel-2 quality bitmasks over-masked water and under-masked cloud shadows. We used the s2cloudless method (Zupanc 2017), which was much more accurate, but cloud shadows still confounded our AWEI thresholding method and necessitated using median imagery to handle shadow outlier pixels. There is promise for deep learning and artificial intelligence-supported cloud (Zupanc 2017; López-Puigdollers et al. 2021; Anzalone et al. 2024) and burn masking (Knopp et al. 2020; Pinto et al. 2021), and we encourage researchers with computational expertise to explore these methods when applying our ESW technique.

It is important to select an appropriate water mapping method for any given research application. We recommend that researchers apply our ESW mapping process (Fig. 2) and code to projects with small, infrequently filled water sources and in regions without dense canopy cover (which can obscure water from Sentinel-2 sensors), such as seasonal savannas. We also recommend that researchers double-check the masks used against the landscape; our analysis does not include areas with mountains, snow, or ice (Zhai et al. 2015; Y. Zhou et al. 2017); and, as solar power increases across southern Africa, solar farms may become more abundant and create false positives for water (Jain and Jain 2017). Researchers should also ensure that the timings of wet and dry seasons for their study area are accurately identified, as they may be different from those in KAZA. In addition, for riparian regions of seasonal inundation (e.g., SIZ), we recommend against using the maximum water fill raster as a mask on imagery, as this approach is better suited for small waterholes that are not likely to spread beyond their maximum fill. When possible, we recommend hand-selecting time periods of clear imagery, as this will result in the cleanest product; however, for long time spans and for large regions like KAZA, we recommend binning imagery in two- or three-month periods and using strict cloud filtering as a trade-off to reduce the large effort required by hand-selection.

## 5 Conclusions

Climate change is expected to intensify rapidly in southern Africa over the coming decades (Engelbrecht et al. 2015; Masson-Delmotte et al. 2021; Black et al. 2024); the accurate mapping of water resources will therefore be critical to tracking aridification trends and their impacts on humans, livestock, agriculture, and wildlife. Our case study of savanna elephant movements supports previous, smaller-scale research (Boyers et al. 2019; Cain et al. 2012; Chamaillé-Jammes, Fritz, et al. 2007; Naidoo et al. 2020) indicating that models omitting small pools of ephemeral surface water are ignoring a major driver of wildlife movement, as many large mammals rely on small water sources to follow a green wave of vegetation change (Bischof et al. 2012; Naidoo et al. 2014; Makati et al. 2025). For example, human-elephant conflict (HEC) spikes during drought, especially near water sources (Montero-Botey et al. 2024; Kairiza et al. 2025). Further investigation into the inter-annual and seasonal variation in HEC and elephant use of small, dynamic waterholes to access vegetation could assist in potential interventions to avoid HEC. In addition, wildlife interactions with veterinary fences influence disease flow between wildlife and livestock (Osofsky 2019; Rosen et al. 2026); models omitting a key driver of animal movements risk mischaracterizing the disease landscape. Projections of future wildlife movements like these are important for predicting ecosystem health, nutrient flow, disease risk, and human-wildlife conflict under climate change, and we have shown here that ephemeral water is a key piece of the puzzle.

Understanding how animal behavior and ecosystem processes interact with climate change cannot occur without accurate maps of surface water at a temporal and spatial scale that matters to wildlife. We hope that the work provided here will extend water mapping accessibility to more researchers in southern Africa, and that our code and methods offer a starting point for water mapping efforts outside of the KAZA Transfrontier Conservation Area.

## Acknowledgements

The authors would like to thank the teams at Ecoexist Trust and the Ministry of Environment, Forestry, and Tourism in Namibia for their work collecting elephant GPS data; Steve Osofsky and Shirley Atkinson for their comments on the manuscript; and WWF-US and the Cornell Atkinson Center for Sustainability for their support of MES. Feedback from Summer Smentek was invaluable in testing our ESW product beyond KAZA, especially on black cotton soils and near solar panel installations. No part of this manuscript was prepared using generative AI tools. We would like to dedicate this paper in memory of Anastacia Makati, whose humor, kindness, and scientific insight will be missed by humans and elephants alike.

## Author Contributions (CRediT)

**MES*:** Conceptualization, Methodology, Software, Validation, Formal Analysis, Writing – Original Draft, Visualization; **AS, GM, PB:** Investigation, Resources, Data Curation, Writing – Review & Editing, Project administration; **RN:** Conceptualization, Investigation, Data Curation, Writing – Review & Editing, Supervision, Funding acquisition.

## Open Research Statement

All surface water raster data are hosted on HydroShare: http://www.hydroshare.org/resource/8d974079c9f146cfa25aaeeab7a8a342. This submission uses novel code; code for Google Earth Engine water mapping are shared here: https://www.hydroshare.org/resource/1eda26a154c34f2c9aa3e2eb41877ad4/ while code for the seasonal fill analysis and elephant water use case study are shared on GitHub: https://github.com/margaret-swift/kaza-water-case-study/. Individual savanna elephant GPS collar data are sensitive due to poaching concerns and cannot be provided publicly.

