## Appendix S1 for "Mapping small-scale ephemeral surface water to inform transfrontier conservation planning in southern Africa"

*Ecosphere: ECS26-0338*

**Mapping small-scale ephemeral surface water to inform transfrontier conservation planning in southern Africa**

Margaret E. Swift*, Anna Songhurst, Graham McCulloch, Piet Beytell, Robin Naidoo

**Contents: Figures and Tables**


| Page | Description |
|---|---|
| **2** | **Figure S1.** Data used in savanna elephant (*Loxodonta africana*) case study |
| **3-9** | **Table S1:** Hand-selected Sentinel-2 imagery dates for SIZ. |
| **10** | **Table S2:** Sentinel-2 image masking data sources and processing. |
| **11** | **Table S3:** Number of validation points for each region, year, and classification. |
| **12** | **Table S4**: Validation results for ESW with and without MWF positive masking. |
| **13** | **Table S5**: Kruskall-Wallis test for pentad differences in proportion of Maximum Water Fill (*pMWF*) |

**Figure S1.** Data used in savanna elephant (*Loxodonta africana*) case study. GPS telemetry data (a) are from 6 male and 6 female elephant collared in Botswana in 2015, 2017, and 2021, and 2 male and 13 female elephant collared in Namibia in 2010, 2016, 2017, and 2020. The case study area of interest at the northeast border of Namibia and Botswana (b) covers a range of savanna vegetation (NDVI in green) near the Kwando River (blue). The Ephemeral Surface Water (ESW) product is mapped in purple, and a veterinary fence crosses the savanna (dashed line).

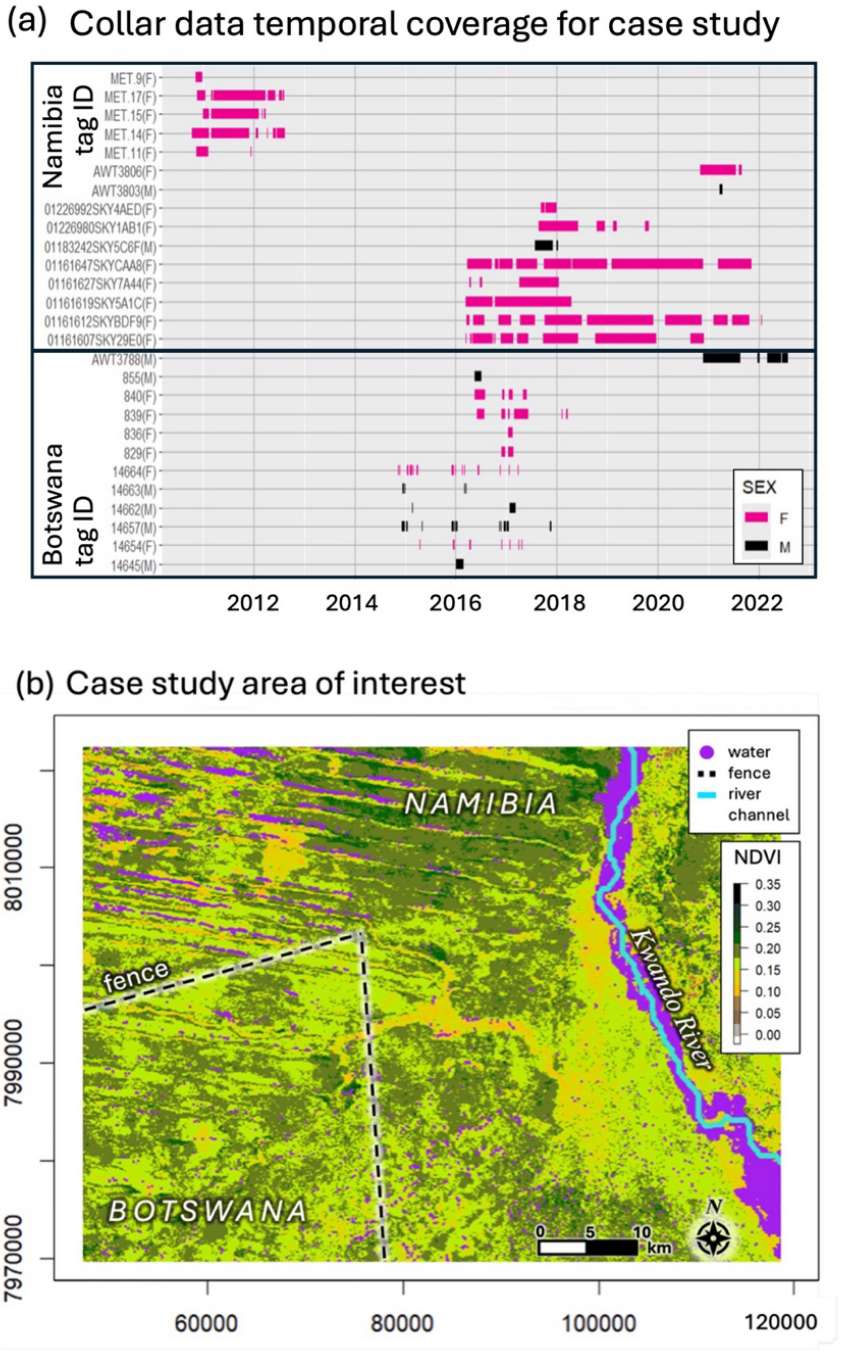

**Table S1:** Hand-selected Sentinel-2 imagery dates for SIZ.

| ID | START | END | NAME | CLOUD_FILTER | CLOUD_PROB |
|---|---|---|---|---|---|
| Zambezi | 12/1/18 | 2/1/19 | 2018_DEC_FEB | 80 | 50 |
| Zambezi | 3/1/19 | 4/30/19 | 2019_MAR_APR | 80 | 50 |
| Zambezi | 5/1/19 | 6/30/19 | 2019_MAY_JUN | 80 | 50 |
| Zambezi | 7/1/19 | 8/31/19 | 2019_JUL_AUG | 80 | 50 |
| Zambezi | 9/1/19 | 11/15/19 | 2019_SEP_NOV | 80 | 50 |
| Zambezi | 11/15/19 | 1/15/20 | 2019_NOV_JAN | 80 | 50 |
| Zambezi | 1/1/20 | 3/1/20 | 2020_JAN_FEB | 80 | 50 |
| Zambezi | 2/1/20 | 3/15/20 | 2020_FEB | 80 | 50 |
| Zambezi | 3/15/20 | 3/31/20 | 2020_MAR | 80 | 50 |
| Zambezi | 3/31/20 | 4/30/20 | 2020_APR | 80 | 50 |
| Zambezi | 5/1/20 | 5/31/20 | 2020_MAY | 80 | 50 |
| Zambezi | 6/1/20 | 6/30/20 | 2020_JUN | 80 | 50 |
| Zambezi | 7/1/20 | 7/31/20 | 2020_JUL | 80 | 50 |
| Zambezi | 8/1/20 | 8/31/20 | 2020_AUG | 80 | 50 |
| Zambezi | 9/1/20 | 9/30/20 | 2020_SEP | 80 | 50 |
| Zambezi | 10/1/20 | 10/31/20 | 2020_OCT | 80 | 50 |
| Zambezi | 11/1/20 | 1/31/21 | 2020_NOV_JAN | 80 | 50 |
| Zambezi | 2/1/21 | 3/31/21 | 2021_FEB_MAR | 80 | 50 |
| Zambezi | 4/1/21 | 4/15/21 | 2021_APR1 | 80 | 50 |
| Zambezi | 4/15/21 | 4/30/21 | 2021_APR2 | 80 | 50 |
| Zambezi | 5/1/21 | 5/31/21 | 2021_MAY | 80 | 50 |
| Zambezi | 6/1/21 | 6/30/21 | 2021_JUN | 80 | 50 |
| Zambezi | 7/1/21 | 7/31/21 | 2021_JUL | 80 | 50 |
| Zambezi | 8/1/21 | 8/31/21 | 2021_AUG | 80 | 50 |
| Zambezi | 9/1/21 | 10/31/21 | 2021_SEP_OCT | 80 | 50 |
| Zambezi | 11/1/21 | 1/31/22 | 2021_NOV_JAN | 80 | 50 |
| Zambezi | 2/1/22 | 2/28/22 | 2022_FEB | 80 | 50 |
| Zambezi | 3/1/22 | 3/31/22 | 2022_MAR | 80 | 50 |
| Zambezi | 4/1/22 | 4/30/22 | 2022_APR | 80 | 50 |
| Zambezi | 5/1/22 | 5/31/22 | 2022_MAY | 80 | 50 |
| Zambezi | 6/1/22 | 6/30/22 | 2022_JUN | 80 | 50 |
| Zambezi | 7/1/22 | 7/31/22 | 2022_JUL | 80 | 50 |
| Zambezi | 8/1/22 | 8/31/22 | 2022_AUG | 80 | 50 |
| Zambezi | 9/1/22 | 9/30/22 | 2022_SEP | 80 | 50 |
| Zambezi | 10/1/22 | 11/30/22 | 2022_OCT_NOV | 80 | 50 |
| Zambezi | 12/1/22 | 12/31/22 | 2022_DEC | 80 | 50 |
| Zambezi | 1/1/23 | 2/15/23 | 2023_JAN_FEB | 80 | 50 |
| Zambezi | 2/15/23 | 3/1/23 | 2023_FEB | 80 | 50 |
| Zambezi | 3/1/23 | 3/31/23 | 2023_MAR | 80 | 50 |
| Zambezi | 4/1/23 | 4/30/23 | 2023_APR | 80 | 50 |
| Zambezi | 5/1/23 | 5/31/23 | 2023_MAY | 80 | 50 |
| Zambezi | 6/1/23 | 6/30/23 | 2023_JUN | 80 | 50 |
| Zambezi | 7/1/23 | 7/31/23 | 2023_JUL | 80 | 50 |
| Zambezi | 8/1/23 | 8/31/23 | 2023_AUG | 80 | 50 |
| Zambezi | 9/1/23 | 11/15/23 | 2023_SEP_OCT | 80 | 50 |
| Zambezi | 10/15/23 | 12/31/23 | 2023_NOV_DEC | 80 | 50 |
| Zambezi | 1/1/24 | 1/31/24 | 2024_JAN | 80 | 50 |
| Zambezi | 2/1/24 | 2/28/24 | 2024_FEB | 80 | 50 |
| Zambezi | 3/1/24 | 4/30/24 | 2024_MAR_APR | 80 | 50 |
| Zambezi | 5/1/24 | 6/30/24 | 2024_MAY_JUN | 80 | 50 |
| Zambezi | 7/1/24 | 8/31/24 | 2024_JUL_AUG | 80 | 50 |
| Zambezi | 9/1/24 | 10/31/24 | 2024_SEP_OCT | 80 | 50 |

| | | | | | |
|---|---|---|---|---|---|
| Zambezi | 10/15/24 | 12/31/24 | 2024_NOV_DEC | 80 | 50 |
| Zambezi | 1/1/25 | 2/15/25 | 2025_JAN | 80 | 50 |
| Zambezi | 2/1/25 | 3/15/25 | 2025_FEB | 80 | 50 |
| Zambezi | 3/15/25 | 3/31/25 | 2025_MAR | 80 | 50 |
| Zambezi | 4/1/25 | 4/30/25 | 2025_APR | 80 | 50 |
| Zambezi | 5/1/25 | 5/31/25 | 2025_MAY | 80 | 50 |
| Zambezi | 6/1/25 | 6/30/25 | 2025_JUN | 80 | 50 |
| Zambezi | 7/1/25 | 7/31/25 | 2025_JUL | 80 | 50 |
| Zambezi | 8/1/25 | 8/31/25 | 2025_AUG | 80 | 50 |
| Zambezi | 9/1/25 | 9/30/25 | 2025_SEP | 80 | 50 |
| Okavango | 1/1/19 | 2/28/19 | 2019_JAN_FEB | 80 | 50 |
| Okavango | 2/15/19 | 3/30/19 | 2019_MAR | 80 | 50 |
| Okavango | 4/1/19 | 5/30/19 | 2019_APR_MAY | 80 | 50 |
| Okavango | 6/1/19 | 7/31/19 | 2019_JUN_JUL | 80 | 50 |
| Okavango | 7/15/19 | 9/15/19 | 2019_AUG_SEP | 80 | 50 |
| Okavango | 9/1/19 | 10/31/19 | 2019_SEP_OCT | 80 | 50 |
| Okavango | 11/1/19 | 12/31/19 | 2019_NOV_DEC | 80 | 50 |
| Okavango | 12/15/19 | 2/28/20 | 2020_DEC_FEB | 80 | 50 |
| Okavango | 3/1/20 | 3/31/20 | 2020_MAR | 80 | 50 |
| Okavango | 4/1/20 | 4/30/20 | 2020_APR | 80 | 50 |
| Okavango | 5/1/20 | 5/31/20 | 2020_MAY | 80 | 50 |
| Okavango | 6/1/20 | 6/30/20 | 2020_JUN | 80 | 50 |
| Okavango | 7/1/20 | 7/31/20 | 2020_JUL | 80 | 50 |
| Okavango | 8/1/20 | 8/31/20 | 2020_AUG | 80 | 50 |
| Okavango | 9/1/20 | 9/30/20 | 2020_SEP | 80 | 50 |
| Okavango | 10/1/20 | 10/31/20 | 2020_OCT | 80 | 50 |
| Okavango | 11/1/20 | 11/30/20 | 2020_NOV | 80 | 50 |
| Okavango | 12/1/20 | 12/31/20 | 2020_DEC | 80 | 50 |
| Okavango | 1/1/21 | 3/15/21 | 2021_JAN_FEB | 80 | 50 |
| Okavango | 3/15/21 | 3/31/21 | 2021_MAR | 80 | 50 |
| Okavango | 4/1/21 | 4/30/21 | 2021_APR | 80 | 50 |
| Okavango | 5/1/21 | 5/31/21 | 2021_MAY | 80 | 50 |
| Okavango | 6/1/21 | 6/30/21 | 2021_JUN | 80 | 50 |
| Okavango | 7/1/21 | 8/31/21 | 2021_JUL_AUG | 80 | 50 |
| Okavango | 9/1/21 | 11/30/21 | 2021_SEP_NOV | 80 | 50 |
| Okavango | 11/1/21 | 1/15/22 | 2021_NOV_JAN | 80 | 50 |
| Okavango | 1/15/22 | 2/15/22 | 2022_JAN_FEB | 80 | 50 |
| Okavango | 2/15/22 | 3/31/22 | 2022_FEB_MAR | 80 | 50 |
| Okavango | 4/1/22 | 5/31/22 | 2022_APR_MAY | 80 | 50 |
| Okavango | 6/1/22 | 7/31/22 | 2022_JUN_JUL | 80 | 50 |
| Okavango | 8/1/22 | 10/31/22 | 2022_AUG_OCT | 80 | 50 |
| Okavango | 11/1/22 | 12/31/22 | 2022_NOV_DEC | 80 | 50 |
| Okavango | 1/1/23 | 3/30/23 | 2023_JAN_MAR | 80 | 50 |
| Okavango | 4/1/23 | 5/30/23 | 2023_APR_MAY | 80 | 50 |
| Okavango | 5/15/23 | 7/31/23 | 2023_JUN_JUL | 80 | 50 |
| Okavango | 8/1/23 | 9/30/23 | 2023_AUG_SEP | 80 | 50 |
| Okavango | 10/1/23 | 11/30/23 | 2023_OCT_NOV | 80 | 50 |
| Okavango | 12/1/23 | 2/15/24 | 2023_DEC_JAN | 80 | 50 |
| Okavango | 2/15/24 | 3/15/24 | 2024_FEB_MAR | 80 | 50 |
| Okavango | 3/15/24 | 4/30/24 | 2024_MAR_APR | 80 | 50 |
| Okavango | 5/1/24 | 5/31/24 | 2024_MAY | 80 | 50 |
| Okavango | 6/1/24 | 6/30/24 | 2024_JUN | 80 | 50 |
| Okavango | 7/1/24 | 7/31/24 | 2024_JUL | 80 | 50 |
| Okavango | 8/1/24 | 9/30/24 | 2024_AUG_SEP | 80 | 50 |
| Okavango | 10/1/24 | 10/31/24 | 2024_OCT | 80 | 50 |

| | | | | | |
|---|---|---|---|---|---|
| Okavango | 10/15/24 | 12/31/24 | 2024_NOV_DEC | 80 | 50 |
| Okavango | 1/1/25 | 2/28/25 | 2025_JAN_FEB | 80 | 50 |
| Okavango | 3/1/25 | 3/31/25 | 2025_MAR | 80 | 50 |
| Okavango | 4/1/25 | 4/30/25 | 2025_APR | 80 | 50 |
| Okavango | 5/1/25 | 5/31/25 | 2025_MAY | 80 | 50 |
| Okavango | 6/1/25 | 6/30/25 | 2025_JUN | 80 | 50 |
| Okavango | 7/1/25 | 8/31/25 | 2025_JUL_AUG | 80 | 50 |
| Okavango | 9/1/25 | 10/15/25 | 2025_SEP | 80 | 50 |
| Makgadikgadi | 1/1/19 | 1/31/19 | 2019_JAN | 30 | 90 |
| Makgadikgadi | 2/1/19 | 2/28/19 | 2019_FEB | 30 | 90 |
| Makgadikgadi | 3/1/19 | 3/31/19 | 2019_MAR | 30 | 90 |
| Makgadikgadi | 4/1/19 | 4/30/19 | 2019_APR | 30 | 90 |
| Makgadikgadi | 5/1/19 | 5/31/19 | 2019_MAY | 30 | 90 |
| Makgadikgadi | 6/1/19 | 6/30/19 | 2019_JUN | 30 | 90 |
| Makgadikgadi | 7/1/19 | 7/31/19 | 2019_JUL | 30 | 90 |
| Makgadikgadi | 8/1/19 | 8/31/19 | 2019_AUG | 30 | 90 |
| Makgadikgadi | 9/1/19 | 9/30/19 | 2019_SEP | 30 | 90 |
| Makgadikgadi | 10/1/19 | 10/31/19 | 2019_OCT | 30 | 90 |
| Makgadikgadi | 11/1/19 | 11/30/19 | 2019_NOV | 30 | 90 |
| Makgadikgadi | 12/1/19 | 12/31/19 | 2019_DEC | 30 | 90 |
| Makgadikgadi | 1/1/20 | 2/28/20 | 2020_JAN_FEB | 30 | 90 |
| Makgadikgadi | 3/1/20 | 4/30/20 | 2020_MAR_APR | 30 | 90 |
| Makgadikgadi | 5/1/20 | 5/31/20 | 2020_MAY | 30 | 90 |
| Makgadikgadi | 6/1/20 | 6/30/20 | 2020_JUN | 30 | 90 |
| Makgadikgadi | 7/1/20 | 7/31/20 | 2020_JUL | 30 | 90 |
| Makgadikgadi | 8/1/20 | 8/31/20 | 2020_AUG | 30 | 90 |
| Makgadikgadi | 9/1/20 | 9/30/20 | 2020_SEP | 30 | 90 |
| Makgadikgadi | 10/1/20 | 10/31/20 | 2020_OCT | 30 | 90 |
| Makgadikgadi | 11/1/20 | 12/31/20 | 2020_NOV_DEC | 30 | 90 |
| Makgadikgadi | 1/1/21 | 3/15/21 | 2021_JAN_FEB | 30 | 90 |
| Makgadikgadi | 3/15/21 | 3/31/21 | 2021_MAR | 30 | 90 |
| Makgadikgadi | 4/1/21 | 4/30/21 | 2021_APR | 30 | 90 |
| Makgadikgadi | 5/1/21 | 5/31/21 | 2021_MAY | 30 | 90 |
| Makgadikgadi | 6/1/21 | 6/30/21 | 2021_JUN | 30 | 90 |
| Makgadikgadi | 7/15/21 | 7/31/21 | 2021_JUL | 30 | 90 |
| Makgadikgadi | 8/1/21 | 8/31/21 | 2021_AUG | 30 | 90 |
| Makgadikgadi | 9/1/21 | 9/30/21 | 2021_SEP | 30 | 90 |
| Makgadikgadi | 10/1/21 | 10/31/21 | 2021_OCT | 30 | 90 |
| Makgadikgadi | 11/1/21 | 12/31/21 | 2021_NOV_DEC | 30 | 90 |
| Makgadikgadi | 1/1/22 | 1/30/22 | 2021_JAN | 30 | 90 |
| Makgadikgadi | 2/1/22 | 2/28/22 | 2022_FEB | 30 | 90 |
| Makgadikgadi | 3/1/22 | 3/31/22 | 2022_MAR | 30 | 90 |
| Makgadikgadi | 4/1/22 | 4/30/22 | 2022_APR | 30 | 90 |
| Makgadikgadi | 5/1/22 | 5/31/22 | 2022_MAY | 30 | 90 |
| Makgadikgadi | 6/1/22 | 6/30/22 | 2022_JUN | 30 | 90 |
| Makgadikgadi | 7/1/22 | 7/31/22 | 2022_JUL | 30 | 90 |
| Makgadikgadi | 8/1/22 | 8/31/22 | 2022_AUG | 30 | 90 |
| Makgadikgadi | 9/1/22 | 10/31/22 | 2022_SEP_OCT | 30 | 90 |
| Makgadikgadi | 11/1/22 | 1/15/23 | 2022_NOV_JAN | 30 | 90 |
| Makgadikgadi | 1/15/23 | 2/28/23 | 2023_JAN_FEB | 30 | 90 |
| Makgadikgadi | 3/1/23 | 3/31/23 | 2023_MAR | 30 | 90 |
| Makgadikgadi | 4/1/23 | 4/30/23 | 2023_APR | 30 | 90 |
| Makgadikgadi | 5/1/23 | 5/31/23 | 2023_MAY | 30 | 90 |
| Makgadikgadi | 6/1/23 | 6/30/23 | 2023_JUN | 30 | 90 |
| Makgadikgadi | 7/1/23 | 8/31/23 | 2023_JUL_AUG | 30 | 90 |

| | | | | | |
|---|---|---|---|---|---|
| Makgadikgadi | 9/1/23 | 9/30/23 | 2023_SEP | 30 | 90 |
| Makgadikgadi | 10/1/23 | 10/31/23 | 2023_OCT | 30 | 90 |
| Makgadikgadi | 11/1/23 | 11/30/23 | 2023_NOV | 30 | 90 |
| Makgadikgadi | 12/1/23 | 12/31/23 | 2023_DEC | 30 | 90 |
| Makgadikgadi | 1/1/24 | 1/31/24 | 2024_JAN | 30 | 90 |
| Makgadikgadi | 2/1/24 | 2/28/24 | 2024_FEB | 30 | 90 |
| Makgadikgadi | 3/1/24 | 3/31/24 | 2024_MAR | 30 | 90 |
| Makgadikgadi | 4/1/24 | 4/30/24 | 2024_APR | 30 | 90 |
| Makgadikgadi | 5/1/24 | 5/31/24 | 2024_MAY | 30 | 90 |
| Makgadikgadi | 6/1/24 | 6/30/24 | 2024_JUN | 30 | 90 |
| Makgadikgadi | 7/1/24 | 7/31/24 | 2024_JUL | 30 | 90 |
| Makgadikgadi | 8/1/24 | 8/31/24 | 2024_AUG | 30 | 90 |
| Makgadikgadi | 9/1/24 | 9/30/24 | 2024_SEP | 30 | 90 |
| Makgadikgadi | 10/1/24 | 10/31/24 | 2024_OCT | 30 | 90 |
| Makgadikgadi | 11/1/24 | 12/31/24 | 2024_NOV_DEC | 30 | 90 |
| Makgadikgadi | 1/1/25 | 1/31/25 | 2025_JAN | 30 | 90 |
| Makgadikgadi | 2/1/25 | 2/28/25 | 2025_FEB | 30 | 90 |
| Makgadikgadi | 3/1/25 | 3/31/25 | 2025_MAR | 30 | 90 |
| Makgadikgadi | 4/1/25 | 4/30/25 | 2025_APR | 30 | 90 |
| Makgadikgadi | 5/1/25 | 5/31/25 | 2025_MAY | 30 | 90 |
| Makgadikgadi | 6/1/25 | 6/30/25 | 2025_JUN | 30 | 90 |
| Makgadikgadi | 7/1/25 | 7/31/25 | 2025_JUL | 30 | 90 |
| Makgadikgadi | 8/1/25 | 8/31/25 | 2025_AUG | 30 | 90 |
| Makgadikgadi | 9/1/25 | 9/30/25 | 2025_SEP | 30 | 90 |
| Linyanti | 1/1/19 | 2/28/19 | 2019_JAN_FEB | 80 | 50 |
| Linyanti | 3/1/19 | 3/31/19 | 2019_MAR-APR | 80 | 50 |
| Linyanti | 5/1/19 | 6/30/19 | 2019_MAY_JUN | 80 | 50 |
| Linyanti | 10/1/19 | 12/31/19 | 2019_OCT_DEC | 80 | 50 |
| Linyanti | 1/1/19 | 2/28/20 | 2020_JAN_FEB | 80 | 50 |
| Linyanti | 3/1/20 | 3/31/20 | 2020_MAR | 80 | 50 |
| Linyanti | 4/1/20 | 4/30/20 | 2020_APR | 80 | 50 |
| Linyanti | 5/1/20 | 5/31/20 | 2020_MAY | 80 | 50 |
| Linyanti | 6/1/20 | 8/31/20 | 2020_JUN_JUL | 80 | 50 |
| Linyanti | 9/1/20 | 10/31/20 | 2020_SEP_OCT | 80 | 50 |
| Linyanti | 11/1/20 | 12/31/20 | 2020_NOV_DEC | 80 | 50 |
| Linyanti | 1/1/21 | 2/28/21 | 2021_JAN_FEB | 80 | 50 |
| Linyanti | 3/1/21 | 3/31/21 | 2021_MAR | 80 | 50 |
| Linyanti | 4/1/21 | 4/30/21 | 2021_APR | 80 | 50 |
| Linyanti | 5/1/21 | 5/31/21 | 2021_MAY | 80 | 50 |
| Linyanti | 5/15/21 | 7/31/21 | 2021_JUN_JUL | 80 | 50 |
| Linyanti | 8/1/21 | 8/31/21 | 2021_AUG | 80 | 50 |
| Linyanti | 9/1/21 | 10/31/21 | 2021_SEP_OCT | 80 | 50 |
| Linyanti | 11/1/21 | 12/31/21 | 2021_NOV_DEC | 80 | 50 |
| Linyanti | 1/1/22 | 2/28/22 | 2022_JAN_FEB | 80 | 50 |
| Linyanti | 3/1/22 | 4/30/22 | 2022_MAR_APR | 80 | 50 |
| Linyanti | 5/1/22 | 5/31/22 | 2022_MAY | 80 | 50 |
| Linyanti | 6/1/22 | 6/30/22 | 2022_JUN | 80 | 50 |
| Linyanti | 7/1/22 | 7/31/22 | 2022_JUL | 80 | 50 |
| Linyanti | 8/1/22 | 9/30/22 | 2022_AUG_SEP | 80 | 50 |
| Linyanti | 10/1/22 | 10/31/22 | 2022_OCT | 80 | 50 |
| Linyanti | 11/1/22 | 11/30/23 | 2022_NOV | 80 | 50 |
| Linyanti | 12/1/22 | 12/31/23 | 2022_DEC | 80 | 50 |
| Linyanti | 1/1/23 | 2/28/23 | 2023_JAN | 80 | 50 |
| Linyanti | 3/1/23 | 3/31/23 | 2023_MAR | 80 | 50 |
| Linyanti | 4/1/23 | 4/30/23 | 2023_APR | 80 | 50 |

| | | | | | |
|---|---|---|---|---|---|
| Linyanti | 5/1/23 | 5/15/23 | 2023_MAY | 80 | 50 |
| Linyanti | 5/15/23 | 6/30/23 | 2023_JUN | 80 | 50 |
| Linyanti | 7/1/23 | 9/30/23 | 2023_JUL_AUG_SEP | 80 | 50 |
| Linyanti | 10/1/23 | 11/30/23 | 2023_OCT_NOV | 80 | 50 |
| Linyanti | 12/1/23 | 1/31/24 | 2024_DEC_JAN | 80 | 50 |
| Linyanti | 2/1/24 | 2/28/24 | 2024_FEB | 80 | 50 |
| Linyanti | 3/1/24 | 4/30/24 | 2024_MAR_APR | 80 | 50 |
| Linyanti | 5/1/24 | 6/30/24 | 2024_MAY_JUN | 80 | 50 |
| Linyanti | 7/1/24 | 8/31/24 | 2024_JUL_AUG | 80 | 50 |
| Linyanti | 9/1/24 | 10/31/24 | 2024_SEP_OCT | 80 | 50 |
| Linyanti | 11/1/24 | 12/31/24 | 2024_NOV_DEC | 80 | 50 |
| Linyanti | 1/1/25 | 2/28/25 | 2025_JAN_FEB | 80 | 50 |
| Linyanti | 3/1/25 | 3/31/25 | 2025_MAR | 80 | 50 |
| Linyanti | 4/1/25 | 4/30/25 | 2025_APR | 80 | 50 |
| Linyanti | 5/1/25 | 5/31/25 | 2025_MAY | 80 | 50 |
| Linyanti | 6/1/25 | 6/30/25 | 2025_JUN | 80 | 50 |
| Linyanti | 7/1/25 | 7/31/25 | 2025_JUL | 80 | 50 |
| Linyanti | 8/1/25 | 9/30/25 | 2025_AUG_SEP | 80 | 50 |
| Chobe | 1/1/19 | 2/28/19 | 2019_JAN_FEB | 80 | 50 |
| Chobe | 3/1/19 | 3/31/19 | 2019_MAR-APR | 80 | 50 |
| Chobe | 5/1/19 | 6/30/19 | 2019_MAY_JUN | 80 | 50 |
| Chobe | 10/1/19 | 12/31/19 | 2019_OCT_DEC | 80 | 50 |
| Chobe | 1/1/19 | 2/28/20 | 2020_JAN_FEB | 80 | 50 |
| Chobe | 4/1/20 | 4/30/20 | 2020_APR | 80 | 50 |
| Chobe | 5/1/20 | 5/31/20 | 2020_MAY | 80 | 50 |
| Chobe | 6/1/20 | 8/31/20 | 2020_JUN_JUL | 80 | 50 |
| Chobe | 9/1/20 | 11/15/20 | 2020_SEP_OCT | 80 | 50 |
| Chobe | 11/1/20 | 1/31/21 | 2020_NOV_JAN | 80 | 50 |
| Chobe | 3/1/21 | 3/31/21 | 2021_MAR | 80 | 50 |
| Chobe | 4/1/21 | 4/30/21 | 2021_APR | 80 | 50 |
| Chobe | 5/1/21 | 6/30/21 | 2021_MAY_JUN | 80 | 50 |
| Chobe | 7/1/21 | 8/31/21 | 2021_JUL_AUG | 80 | 50 |
| Chobe | 9/1/21 | 11/30/21 | 2021_SEP_NOV | 80 | 50 |
| Chobe | 12/1/21 | 1/31/22 | 2021_DEC_JAN | 80 | 50 |
| Chobe | 1/1/22 | 2/28/22 | 2022_JAN_FEB | 80 | 50 |
| Chobe | 3/1/22 | 4/15/22 | 2022_MAR_APR | 80 | 50 |
| Chobe | 4/15/22 | 5/31/22 | 2022_MAY | 80 | 50 |
| Chobe | 9/1/22 | 10/31/22 | 2022_SEP_OCT | 80 | 50 |
| Chobe | 11/1/22 | 12/31/22 | 2022_NOV | 80 | 50 |
| Chobe | 2/1/23 | 3/31/23 | 2023_FEB_MAR | 80 | 50 |
| Chobe | 4/1/23 | 4/30/23 | 2023_APR | 80 | 50 |
| Chobe | 5/1/23 | 7/30/23 | 2023_MAY_JUL | 80 | 50 |
| Chobe | 8/1/23 | 9/30/23 | 2023_AUG_SEP | 80 | 50 |
| Chobe | 10/1/23 | 11/30/23 | 2023_OCT_NOV | 80 | 50 |
| Chobe | 1/1/24 | 2/28/24 | 2024_JAN_FEB | 80 | 50 |
| Chobe | 3/1/24 | 3/31/24 | 2024_MAR | 80 | 50 |
| Chobe | 4/1/24 | 4/30/24 | 2024_APR | 80 | 50 |
| Chobe | 5/1/24 | 5/31/24 | 2024_MAY | 80 | 50 |
| Chobe | 6/1/24 | 7/31/24 | 2024_JUN_JUL | 80 | 50 |
| Chobe | 8/1/24 | 8/31/24 | 2024_AUG | 80 | 50 |
| Chobe | 9/1/24 | 10/31/24 | 2024_SEP_OCT | 80 | 50 |
| Chobe | 11/1/24 | 1/31/25 | 2025_NOV_JAN | 80 | 50 |
| Chobe | 2/1/25 | 3/15/25 | 2025_FEB | 80 | 50 |
| Chobe | 3/15/25 | 3/31/25 | 2025_MAR | 80 | 50 |
| Chobe | 4/1/25 | 4/30/25 | 2025_APR | 80 | 50 |

| | | | | | |
|---|---|---|---|---|---|
| Chobe | 5/1/25 | 5/31/25 | 2025_MAY | 80 | 50 |
| Chobe | 6/1/25 | 6/15/25 | 2025_JUN | 80 | 50 |
| Chobe | 6/15/25 | 8/30/25 | 2025_JUL_AUG | 80 | 50 |
| Chobe | 9/1/25 | 9/30/25 | 2025_SEP | 80 | 50 |
| Chobe | 10/1/25 | 10/20/25 | 2025_OCT | 80 | 50 |
| Kariba | 1/1/19 | 2/28/19 | 2019_JAN_FEB | 80 | 50 |
| Kariba | 3/1/19 | 3/31/19 | 2019_MAR-APR | 80 | 50 |
| Kariba | 5/1/19 | 6/30/19 | 2019_MAY_JUN | 80 | 50 |
| Kariba | 7/1/19 | 8/31/19 | 2019_JUL_AUG | 80 | 50 |
| Kariba | 9/1/19 | 9/30/19 | 2019_SEP | 80 | 50 |
| Kariba | 10/1/19 | 10/31/19 | 2019_OCT | 80 | 50 |
| Kariba | 11/1/19 | 11/30/19 | 2019_NOV | 80 | 50 |
| Kariba | 12/1/19 | 12/31/19 | 2019_DEC | 80 | 50 |
| Kariba | 1/1/20 | 2/28/20 | 2020_JAN_FEB | 80 | 50 |
| Kariba | 3/1/20 | 3/31/20 | 2020_MAR | 80 | 50 |
| Kariba | 4/1/20 | 4/30/20 | 2020_APR | 80 | 50 |
| Kariba | 5/1/20 | 5/31/20 | 2020_MAY | 80 | 50 |
| Kariba | 6/1/20 | 8/31/20 | 2020_JUN_JUL | 80 | 50 |
| Kariba | 9/1/20 | 11/15/20 | 2020_SEP_OCT | 80 | 50 |
| Kariba | 11/1/20 | 12/31/20 | 2020_NOV_DEC | 80 | 50 |
| Kariba | 1/1/21 | 3/15/21 | 2021_JAN_MAR | 80 | 50 |
| Kariba | 3/15/21 | 4/30/21 | 2021_MAR_APR | 80 | 50 |
| Kariba | 5/1/21 | 6/30/21 | 2021_MAY_JUN | 80 | 50 |
| Kariba | 7/1/21 | 8/31/21 | 2021_JUL_AUG | 80 | 50 |
| Kariba | 9/1/21 | 10/31/21 | 2021_SEP_OCT | 80 | 50 |
| Kariba | 11/1/21 | 12/31/21 | 2021_NOV_DEC | 80 | 50 |
| Kariba | 1/1/22 | 2/28/22 | 2022_JAN_FEB | 80 | 50 |
| Kariba | 3/1/22 | 4/15/22 | 2022_MAR_APR | 80 | 50 |
| Kariba | 4/15/22 | 5/31/22 | 2022_MAY | 80 | 50 |
| Kariba | 6/1/22 | 7/31/22 | 2022_JUN_JUL | 80 | 50 |
| Kariba | 8/1/22 | 8/31/22 | 2022_AUG | 80 | 50 |
| Kariba | 9/1/22 | 10/31/22 | 2022_SEP_OCT | 80 | 50 |
| Kariba | 11/1/22 | 12/31/22 | 2022_NOV_DEC | 80 | 50 |
| Kariba | 1/1/23 | 2/28/23 | 2023_JAN_FEB | 80 | 50 |
| Kariba | 3/1/23 | 3/31/23 | 2023_MAR | 80 | 50 |
| Kariba | 4/1/23 | 4/30/23 | 2023_APR | 80 | 50 |
| Kariba | 5/1/23 | 5/31/23 | 2023_MAY | 80 | 50 |
| Kariba | 6/1/23 | 6/30/23 | 2023_JUN | 80 | 50 |
| Kariba | 7/1/23 | 7/31/23 | 2023_JUL | 80 | 50 |
| Kariba | 8/1/23 | 9/30/23 | 2023_AUG_SEP | 80 | 50 |
| Kariba | 10/1/23 | 11/30/23 | 2023_OCT_NOV | 80 | 50 |
| Kariba | 12/1/23 | 1/31/24 | 2023_DEC_JAN | 80 | 50 |
| Kariba | 2/1/24 | 2/28/24 | 2024_FEB | 80 | 50 |
| Kariba | 3/1/24 | 3/31/24 | 2024_MAR | 80 | 50 |
| Kariba | 4/1/24 | 4/30/24 | 2024_APR | 80 | 50 |
| Kariba | 5/1/24 | 5/30/24 | 2024_MAY | 80 | 50 |
| Kariba | 6/1/24 | 6/30/24 | 2024_JUN | 80 | 50 |
| Kariba | 7/1/24 | 7/31/24 | 2024_JUL | 80 | 50 |
| Kariba | 8/1/24 | 9/30/24 | 2024_AUG_SEP | 80 | 50 |
| Kariba | 10/1/24 | 10/31/24 | 2024_OCT | 80 | 50 |
| Kariba | 11/1/24 | 12/31/24 | 2025_NOV_DEC | 80 | 50 |
| Kariba | 1/1/25 | 2/28/25 | 2025_JAN_FEB | 80 | 50 |
| Kariba | 3/1/25 | 3/31/25 | 2025_MAR | 80 | 50 |
| Kariba | 4/1/25 | 4/30/25 | 2025_APR | 80 | 50 |
| Kariba | 5/1/25 | 5/31/25 | 2025_MAY | 80 | 50 |

| | | | | | |
|---|---|---|---|---|---|
| Kariba | 6/1/25 | 6/15/25 | 2025_JUN | 80 | 50 |
| Kariba | 6/15/25 | 8/30/25 | 2025_JUL_AUG | 80 | 50 |
| Kariba | 9/1/25 | 9/30/25 | 2025_SEP | 80 | 50 |
| Kariba | 10/1/25 | 10/31/25 | 2025_OCT | 80 | 50 |
| Kariba | 11/1/25 | 11/30/25 | 2025_NOV | 80 | 50 |

**Table S2:** Sentinel-2 image masking data sources and processing.

| **Mask Type** | **Feature** | **Resolution / type** | **Source** | **Processing** |
|---|---|---|---|---|
| negative | clouds | 10m | Sentinel-2 bands | s2Cloudless method |
| negative | roads | vector | OpenStreetMap | Buffer by 5m |
| negative | settled areas | 10m | World Settlement Footprint 2015 | Masked as is |
| negative | buildings | 4m | Google Open Buildings 2.5D Temporal Dataset | >50% confidence |
| negative | burn scarring | 250m | MODIS MOD44W V6, Burned Area bitmask (MCD64A1) | Removed any pixels marked as ‘burned’ within the last two months |
| positive | Permanent surface water | 30m | JRC Global Surface Water Product, Pekel et al 2016 | Masked as is |

**Table S3:** Number of validation points for each region, year, and classification.

| | Wet season | | Dry season | |
|---|---|---|---|---|
| **REGION** | Not water | Not water | Water | Water |
| A (Kalala) | 320 | 576 | 268 | 314 |
| B (Cuando Cubango) | 554 | 260 | 98 | 226 |
| C (Okavango) | 352 | 344 | 224 | 490 |
| D (Chobe River) | 364 | 532 | 286 | 282 |
| E (Zambezi River) | 596 | 266 | 510 | 408 |
| F (Ngamiland) | 312 | 418 | 0 | 396 |
| G (Makgadikgadi) | 394 | 196 | 0 | 302 |
| **TOTAL** | **2992** | **2592** | **1386** | **2418** |

**Table S4:** Validation results for ESW with and without MWF positive masking. See Table 1 for a complete comparison of ESW with other methods. Bold values indicate areas where one method outperforms the other by over 2 percentage points.

| | | ESW with MWF | | | | ESW without MWF | | | |
|---|---|---|---|---|---|---|---|---|---|
| | | July 2019 (dry) | | March 2021 (wet) | | July 2019 (dry) | | March 2021 (wet) | |
| **Water Type** | **Region** | *water* | *not water* | *water* | *not water* | *water* | *not water* | *water* | *not water* |
| **Scattered waterholes** | A - Kalala | 94 | 100 | **82** | 100 | **97** | 100 | 74 | 99 |
| | B - Kwando | **61** | 99 | **87** | 98 | 5 | 100 | 41 | 100 |
| **Floodplain** | C - Okavango* | **97** | 98 | 98 | 97 | 81 | 100 | 99 | 90 |
| | D - Chobe | **97** | 100 | 97 | 99 | 14 | 100 | 98 | 100 |
| | E - Zambezi* | **82** | 99 | 99 | **87** | 78 | 100 | 99 | 84 |
| **Seasonally dry** | F - Ngamiland | *NA* | 100 | 87 | 100 | *NA* | 100 | **91** | 100 |
| | G - Makgadikgadi* | *NA* | 100 | 95 | 97 | *NA* | 100 | 92 | 99 |

**Table S5**: Kruskall-Wallis test for pentad differences in proportion of Maximum Water Fill (*pMWF*)

| | Nov - Feb | Feb - May | May - July | July - Sept |
|---|---|---|---|---|
| Feb - May | << **0.01** | | | |
| May - July | 0.25 | << **0.01** | | |
| July - Sept | 1 | << **0.01** | 0.018 | |
| Sept - Nov | 0.056 | << **0.01** | << **0.01** | 0.476 |
